# SARS-CoV-2 nsp15 preferentially degrades AU-rich dsRNA via its dsRNA nickase activity

**DOI:** 10.1101/2023.12.11.571056

**Authors:** Xionglue Wang, Bin Zhu

## Abstract

It has been proposed that coronavirus nsp15 mediates evasion of host cell double-stranded (ds) RNA sensors via its uracil-specific endoribonuclease activity. However, how nsp15 processes viral dsRNA, commonly considered as a genome replication intermediate, remains elusive. Previous research has mainly focused on short single-stranded RNA as substrates, and whether nsp15 prefers single-stranded or double-stranded RNA for cleavage is controversial. In the present work, we prepared numerous RNA substrates, including both long substrates mimicking the viral genome and short defined RNA, to clarify the substrate preference and cleavage pattern of SARS-CoV-2 nsp15. We demonstrated that SARS-CoV-2 nsp15 preferentially cleaved flexible pyrimidine nucleotides located in AU-rich areas and mismatch-containing areas in dsRNA via a nicking manner. The AU content and distribution in dsRNA along with the RNA length affected cleavage by SARS-CoV-2 nsp15. Because coronavirus genomes generally have a high AU content, our work supported the mechanism that coronaviruses evade the antiviral response mediated by host cell dsRNA sensors by using nsp15 dsRNA nickase to directly cleave dsRNA intermediates formed during genome replication and transcription.

## INTRODUCTION

Severe acute respiratory syndrome coronavirus 2 (SARS-CoV-2) is the pathogen responsible for the coronavirus disease 2019 (COVID-19) pandemic (Gorbalenya et al., 2020), which, since emerging in late 2019, has resulted in approximately 700 million infections and seven million deaths worldwide (Worldometer, 2023), severely impacting the global economy. Currently, new strains of SARS-CoV-2 continue to arise (Carabelli et al., 2023), resulting in new cases and recurring infections. At the same time, the issue of long COVID cannot be disregarded (Bowe et al., 2023; Davis et al., 2023).

Coronaviruses are a diverse class of single-stranded (ss) positive-sense RNA viruses with a genome size of approximately 30,000 nucleotides, a fact that makes them the largest of the known RNA viruses (V’Kovski et al., 2021). Coronaviruses encode 15–16 nonstructural proteins (nsp) to facilitate genome replication and transcription. Among these, nsp7–16 are considered key components of the viral replication-transcription complex (RTC), while nsp12–16 are primarily involved in manipulating the viral RNA (Snijder et al., 2016). Nevertheless, our understanding of these proteins remains inadequate, and the lack of in-depth biochemical and enzymological studies have impeded our understanding of the molecular mechanism of coronaviral genome replication and transcription. Nsp15 is one of the nsps for which the function and exact role in genome replication and transcription remain elusive.

Coronavirus nsp15 was reported as a hexamer with every protomer comprising three domains: the N-terminal domain, which plays a key role in stabilizing the hexamer; the middle domain; and the C-terminal domain, which belongs to a uracil-specific endoribonuclease (EndoU) family (Bhardwaj et al., 2008; Guarino et al., 2005; Joseph et al., 2007; Kim et al., 2020; Pillon et al., 2021; Ricagno et al., 2006). The EndoU domain is found in all kingdoms of life (Kim et al., 2020), and in RNA viruses, it is chiefly found in nidoviruses. However, not all nidoviruses have an EndoU domain; only those infecting vertebrates, such as coronaviruses and arteriviruses, have evolved EndoU domains (Deng and Baker, 2018). One speculation about this evolutionary phenomenon is that vertebrate nidoviruses evolved EndoU domains in order to counteract the vertebrate-specific innate immune responses against viral RNA, such as the interferon system. This hypothesis is primarily based on several studies that demonstrated that the EndoU activity of coronavirus nsp15 did not directly affect coronavirus replication (Deng et al., 2017; Kang et al., 2007; Kindler et al., 2017), but instead suppressed the antiviral response of the host cell (Deng et al., 2017; Deng et al., 2019; Kindler et al., 2017; Otter et al., 2023; Wu et al., 2020). It was suggested that the EndoU activity of coronavirus nsp15 mediates evasion of host cell double-stranded (ds) RNA sensors by reducing dsRNA produced by viral genome replication and transcription (Deng et al., 2017; Kindler et al., 2017), and two explanations have been proposed to clarify this process. A study by Hackbart M et al. (Hackbart et al., 2020) concluded that coronavirus nsp15 uses its EndoU activity to limit the abundance and length of the 5’-polyU of the viral negative-sense strand RNA, which can fold back and generate stem-loop structures by hybridizing with an A/G-rich domain on the negative-sense strand RNA. This stem-loop structure may be recognized as dsRNA by host cell dsRNA sensors. A study by Ancar R et al. (Ancar et al., 2020) concluded that coronavirus nsp15 used its EndoU activity to cleave U↓A and C↓A sequences within the viral positive-sense strand RNA, including the pre-3’-polyA site, thereby inhibiting viral negative-sense strand RNA synthesis and dsRNA accumulation. Although dsRNA intermediates formed during viral genome replication and transcription are widely acknowledged to be the most probable pathogen-associated molecular patterns (PAMPs) to activate host cell dsRNA sensors (Chen and Hur, 2022; Deng et al., 2017; Hur, 2019; Kindler et al., 2017; Weber et al., 2006), it is unclear whether coronavirus nsp15 directly cleaves such intermediates. One study examining the biochemical properties of coronavirus nsp15 found greater efficacy in the cleavage of dsRNA than ssRNA (Ivanov et al., 2004), while other studies reported that coronavirus nsp15 was more efficient at cleaving ssRNA than dsRNA (Bhardwaj et al., 2004; Frazier et al., 2022). Furthermore, it has been found that when ssRNA forms secondary structures, nsp15 shows a preference for cleaving more flexible Us, particularly those located on loops (Bhardwaj et al., 2006; Salukhe et al., 2023). However, studies of the SARS-CoV-2 nsp15 structure have revealed specific interactions between nsp15 hexamers and dsRNA (Frazier et al., 2022; Ito et al., 2023; Perry et al., 2021), but no significant interactions between nsp15 hexamers and ssRNA, except for the active site pocket (Frazier et al., 2021; Pillon et al., 2021). This discrepancy existing between coronavirus nsp15 biochemical properties and structural studies—as well as our interest in the mechanism of nsp15-mediated immune evasion—motivated us to further investigate the coronavirus nsp15 substrate preference, and therefore in this study, we focused on SARS-CoV-2 nsp15. In previous work, the RNA substrates tested were limited, which may hinder the full characterization of nsp15 activity. In the present work, we prepared a large variety of RNA substrates, including both long substrates mimicking the viral genome and short defined RNA, to clarify the substrate preference and cleavage pattern of SARS-CoV-2 nsp15.

We initially created RNA substrates to mimic ssRNA and dsRNA present during SARS-CoV-2 genome replication and transcription, and found that with RNA substrates containing a high AU content, nsp15 exhibited a significant preference and specificity for cleaving dsRNA, with distinct cleavage sites compared to ssRNA. We then identified that nsp15 preferentially cleaved consecutive Us located in AU-rich areas in dsRNA, and further demonstrated that the relaxed structure of AU-rich areas facilitated U-flipping, making it easier for nsp15 to recognize and cleave the U sites within AU-rich areas. Furthermore, we found that in addition to the AU content and distribution in RNA, the RNA length also affected the substrate preference of nsp15, likely due to differences in the manner by which nsp15 binds to ssRNA and dsRNA. Finally, we clarified that nsp15 cleaved dsRNA as a dsRNA nickase. On the whole, our work supported the mechanism that coronaviruses evade the antiviral response mediated by host cell dsRNA sensors by using nsp15 dsRNA nickase to directly cleave dsRNA intermediates formed during genome replication and transcription.

## RESULTS

### SARS-CoV-2 nsp15 preferentially degrades dsRNA with a high AU content over ssRNA

Most prior studies on the biochemical properties of coronavirus nsp15 employed short RNA substrates ranging from a few to tens of nucleotides, which might not present full recognition elements for nsp15. To closely mimic the naturally occurring RNA during viral genome replication and transcription, we designed an approximately 1-kb chimeric RNA sequence to mimic the SARS-CoV-2 genome (Figure 1A; Table S1) by splicing multiple distinguishing regions from the SARS-CoV-2 genome that were likely to contain potential physiologic targets of nsp15, including the 5’ untranslated region (UTR), 3’ UTR, the core sequence (CS, 5’-ACGAAC-3’) of the transcription regulatory sequence (TRS) (V’Kovski et al., 2021), two sequences located immediately adjacent to open reading frames (ORFs) but lacking the 5’-ACGAAC-3’ sequence, and several complete or incomplete ORF sequences. This chimeric SARS-CoV-2 mini-genome dsRNA was prepared by *in vitro* transcription (IVT) of the positive-sense and negative-sense strand RNA, following by annealing both strands into dsRNA. To prevent interference from dsRNA by-products generated during *in vitro* transcription, we utilized VSW-3 RNA polymerase (Xia et al., 2022), which has previously been demonstrated to effectively reduce these by-products. Furthermore, to exclude the effect of potential nuclease contamination, we purified both the wild-type SARS-CoV-2 nsp15 with a 6× His tag at the N-terminus and its active-site mutant, H234A, as a control for the cleavage reaction, by metal affinity chromatography and gel filtration chromatography (Figure 1B). Previous studies reported that nsp15 required Mn^2+^ for optimal EndoU activity (Bhardwaj et al., 2004; Huang et al., 2023; Ivanov et al., 2004; Zhang et al., 2018); therefore, we initially conducted the cleavage reaction at a Mn^2+^ concentration of 5 mM, which was commonly used in previous studies. We observed that nsp15 efficiently cleaved both ssRNA and dsRNA substrates under this condition (Figures S1A and S1B). However, as 5 mM Mn^2+^ is physiologically irrelevant (Bowman and Aschner, 2014; Chen et al., 2018), we attempted to reduce the concentration of Mn^2+^ to either 0.5 mM or zero while simultaneously increasing the concentration of nsp15 to facilitate cleavage of dsRNA substrates to a similar extent. Surprisingly, nsp15 cleaved the dsRNA substrates significantly more efficiently than the ssRNA substrates under these two conditions (Figures 1C and S1C). To verify the altered substrate preference of nsp15 under different Mn^2+^ concentrations and eliminate the possible effect of nsp15 concentration changes, we conducted the cleavage reaction with the same nsp15 concentration but with various reaction times at various Mn^2+^ concentrations to facilitate cleavage of dsRNA substrates to a similar extent. We found that the cleavage preference of nsp15 for dsRNA was weakened as the Mn^2+^ concentration increased beyond 0.5 mM (Figure S1D). In contrast, no significant change occurred in the cleavage preference of nsp15 for dsRNA when the Mn^2+^ concentration was lower than 0.5 mM (Figure S1E). We then examined the influence of four additional divalent metal ions on the cleavage activity of nsp15 and found that Mg^2+^ and Ca^2+^ also slightly increased the cleavage activity of nsp15, which was in agreement with previous studies (Bhardwaj et al., 2004; Ivanov et al., 2004; Zhang et al., 2018). However, Mg^2+^ and Ca^2+^ did not alter the cleavage preference of nsp15 for dsRNA, whereas Zn^2+^ and Cu^2+^ inhibited the cleavage activity of nsp15, which has not been reported before (Figure 1D). To investigate whether nsp15 had a similar cleavage preference for dsRNA as other sequences, we extracted two sequences from either the SARS-CoV-2, *E. coli*, or *Thermus aquaticus* (*Taq*) genome (Figure 1E; Table S1), and prepared ssRNA and dsRNA substrates by the same method as previously described. Results from the cleavage of these substrates by nsp15 showed a minimal variance in the nsp15 cleavage efficiency of dsRNA substrates derived from the same genome, but a significant difference in the cleavage efficiency of dsRNA substrates derived from different genomes; however, there was no significant difference in the cleavage efficiency of ssRNA substrates derived from different genomes (Figures 1F and S1F). We discovered that one of the most significant differences among various genome-derived RNA substrates consists in their AU content (Figures 1A and 1E), with SARS-CoV-2 RNA possessing a significantly higher AU content (>60%) than that of the RNA from other genomes, signifying that the cleavage preference for dsRNA of nsp15 is positively related to the AU content of RNA substrates.

**Figure 1.**
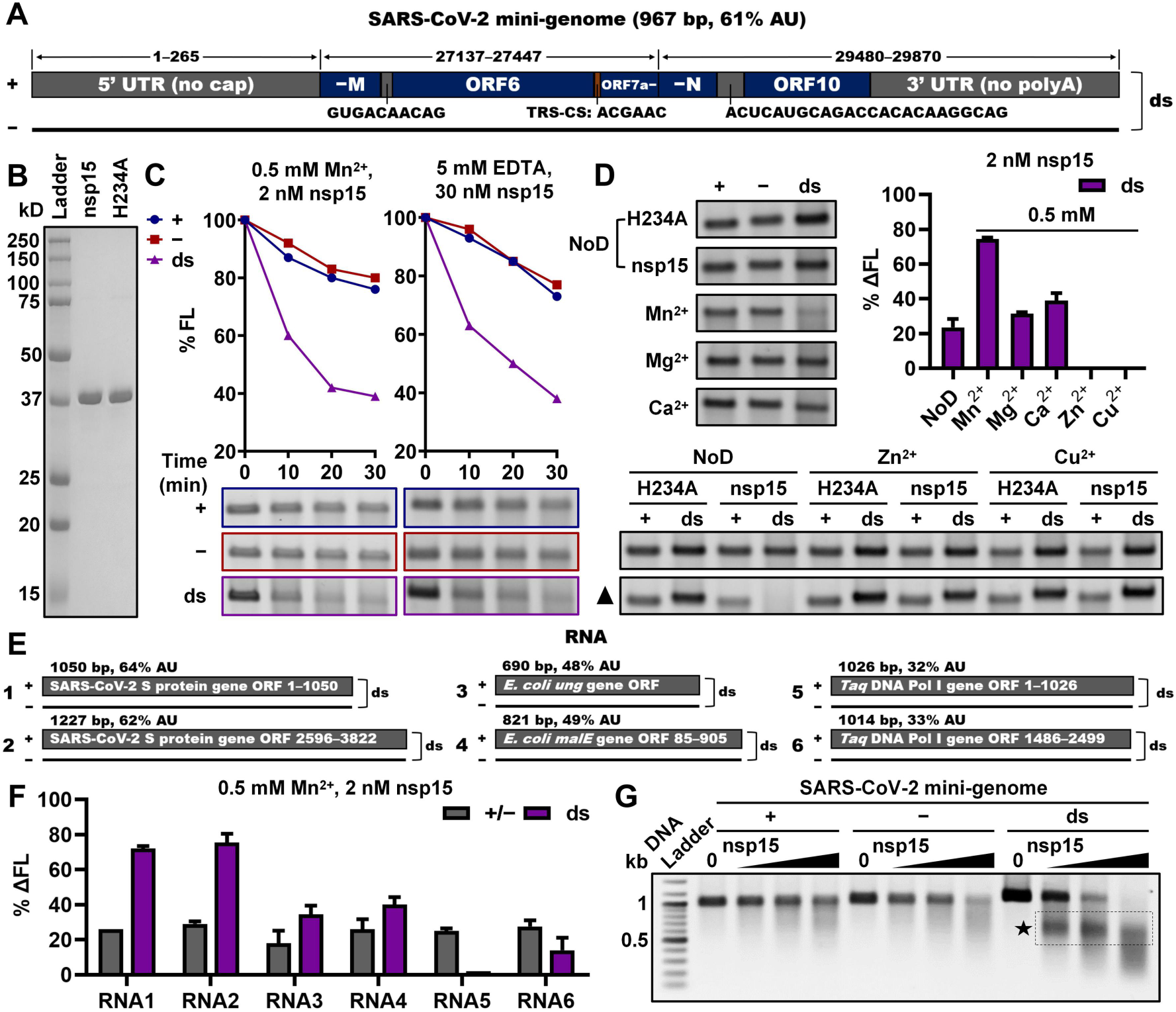
SARS-CoV-2 nsp15 preferentially degrades dsRNA with a high AU content over ssRNA. (**A**) Schematic showing the SARS-CoV-2 mini-genome construction. 1–265, 27,137–27,447, and 29,480–29,870 indicate the corresponding sequence ranges in the SARS-CoV-2 genome. +, −, and ds refer to positive-sense ssRNA, negative-sense ssRNA, and dsRNA, respectively; TRS-CS, the core sequence of the transcription regulatory sequence; 5’ UTR (no cap), 5’ untranslated region without a 5’ cap; −M, right part of the ORF encoding the membrane protein; ORF6, the sixth ORF of the SARS-CoV-2 genome; ORF7a−, left part of the seventh ORF of the SARS-CoV-2 genome; −N, right part of the ORF encoding the nucleocapsid protein; ORF10, the last ORF of the SARS-CoV-2 genome; 3’ UTR (no polyA), 3’ untranslated region without the polyA tail. (**B**) SDS-PAGE analysis of purified wild-type SARS-CoV-2 nsp15 (approximately 40 kDa including an N-terminal 6× His tag) and its active-site mutant, H234A. (**C**) Cleavage of the ssRNA and dsRNA substrates related to the SARS-CoV-2 mini-genome (as shown in **A**) by nsp15 in the presence of 0.5 mM Mn^2+^ or 5 mM EDTA with various reaction times. Remaining full-length RNA substrates after nsp15 cleavage were quantified as % FL. (**D**) Cleavage of the ssRNA and dsRNA substrates related to the SARS-CoV-2 mini-genome by nsp15 in the presence of various divalent metal ions. NoD refers to no ion control. The black triangle indicates increasing the concentration of nsp15 to 30 nM. Reduction of the full-length RNA substrates by nsp15 cleavage was quantified as % ΔFL. (**E**) Schematic representation of RNA substrates 1–6. +, −, and ds refer to positive-sense ssRNA, negative-sense ssRNA, and dsRNA, respectively. (**F**) Cleavage of RNA substrates 1–6 by nsp15 in the presence of 0.5 mM Mn^2+^. Reduction of the full-length RNA substrates by nsp15 cleavage was quantified as % ΔFL. (**G**) Cleavage of the ssRNA and dsRNA substrates related to the SARS-CoV-2 mini-genome by various concentrations of nsp15 (0, 1.5, 5, and 15 nM) in the presence of 0.5 mM Mn^2+^ for 20 min. The prominent gel bands indicating specific cleavage are marked by a black pentagram and a black dashed box. For (**D and F**), reactions containing the same number of nsp15 H234A mutants were used as negative controls. The average and standard deviation for at least two independent reactions are graphed.

Moreover, the gel electrophoresis results of the cleavage products of the SARS-CoV-2 mini-genome dsRNA substrate showed prominent bands (boxed and indicated by black pentagrams in Figures 1G and S1G) with mobilities close to those of the 500-bp DNA marker, whereas no corresponding bands were observed from the cleavage products of related ssRNA substrates (each of the two complementary strands) (Figure 1G), suggesting that nsp15 displays a specific cleavage preference for certain sites in the middle of this dsRNA substrate, but not for the corresponding sites in related ssRNA substrates. Additionally, it was demonstrated that this cleavage preference of nsp15 was not dependent on Mn^2+^ or any other divalent metal ion (Figures S1H and S1I). Given the presence of a TRS-CS in the middle of the SARS-CoV-2 mini-genome RNA (Figure 1A), we investigated whether nsp15 preferentially cleaved this TRS-CS in the form of dsRNA. We prepared a variant of the SARS-CoV-2 mini-genome dsRNA substrate with this TRS-CS deleted (Figure S1J; Table S1) and found that specific cleavage of this variant by nsp15 was also detected, with no significant difference in the cleavage efficiency between this variant and the original SARS-CoV-2 mini-genome dsRNA substrate (Figure S1K), suggesting that the cleavage preference of nsp15 is TRS-CS-independent.

### Identification of the cleavage sites of SARS-CoV-2 nsp15 on the dsRNA substrates derived from the SARS-CoV-2 mini-genome

To locate the preferred cleavage sites of nsp15 on the 967-bp SARS-CoV-2 mini-genome dsRNA (hereinafter O967), we created several variants of this substrate by deleting specific sequences from the original substrate (Figure 2A; Table S1). Based on the size of the fragments after the cleavage of O967 by nsp15, we deduced that the preferred cleavage site was in the middle region of O967. We selected the middle 200 bp of O967 and divided it into left and right halves (Figure 2A, colored in blue and yellow, respectively). First, the middle 200-bp region was deleted from O967 to produce substrate dm200. When dm200 was treated with nsp15, the prominent product band (marked by black pentagrams in Figure 2B) indicating specific cleavage products disappeared. Although certain nonspecific cleavage still occurred, as indicated by the smear in the gel, the overall cleavage efficiency significantly decreased with dm200. However, when the left or right half 100 bp of the middle 200-bp region was deleted, as in the dmL100 and dmR100 substrates, cleavage by nsp15 still produced specific fragments, as revealed by the gel bands indicated by black pentagrams (Figure 2B), suggesting the existence of at least one nsp15 preferred cleavage site within both the left and right halves of this 200-bp region.

**Figure 2.**
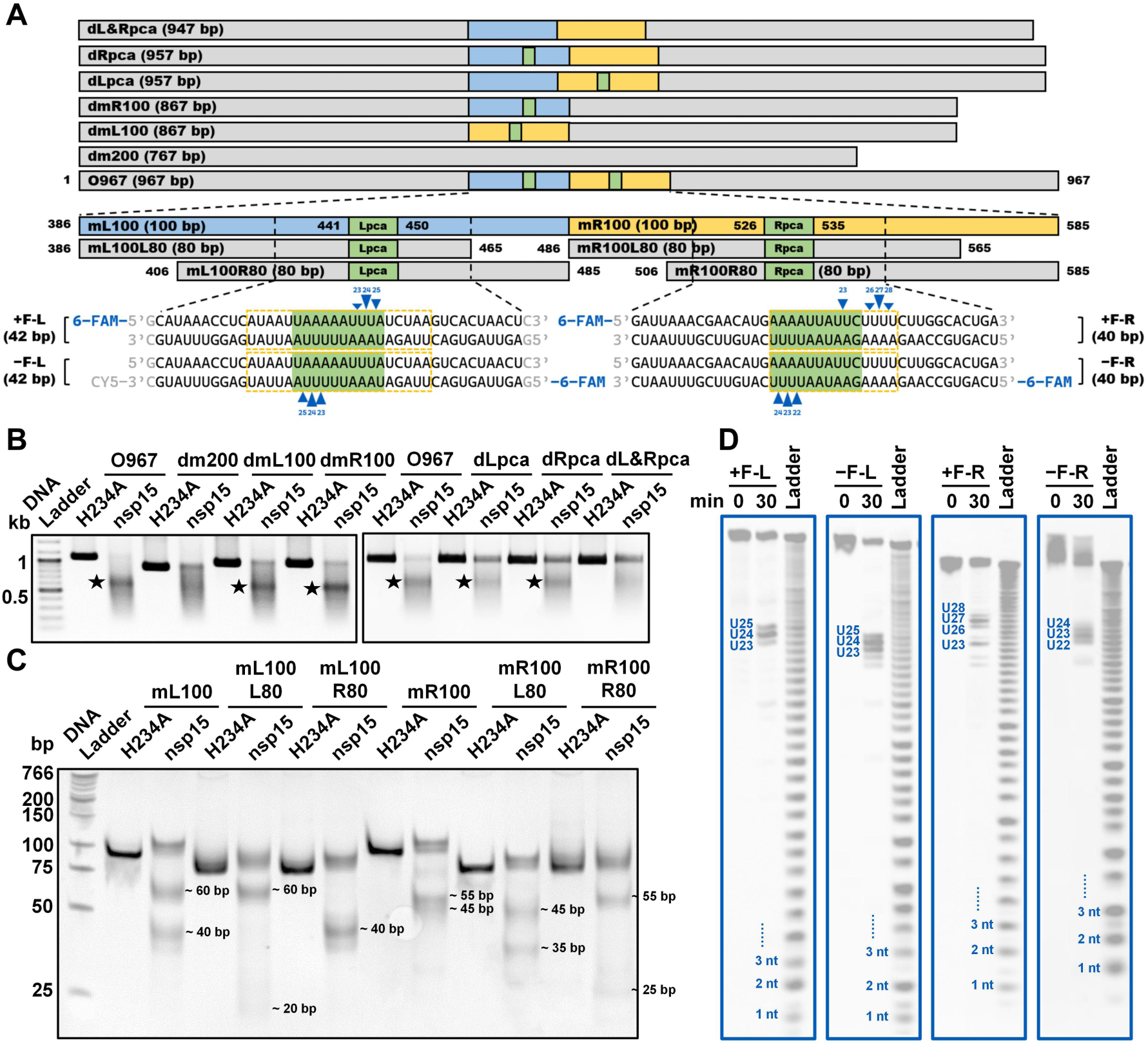
Identification of the cleavage sites of SARS-CoV-2 nsp15 on the dsRNA substrates derived from the SARS-CoV-2 mini-genome. (**A**) Schematic showing the dsRNA substrates derived from the SARS-CoV-2 mini-genome (the original 967-bp dsRNA substrate is designated as O967). Three to four sites identified with the strongest cleavage in every strand of the 6-FAM-labeled dsRNA substrates are marked with blue triangles, and the cleavage efficiency is indicated by the height of the triangle. The AU-rich areas containing these sites are marked by orange dashed boxes. Lpca, left preferred cleavage area; Rpca, right preferred cleavage area. (**B**) Cleavage of the O967 substrate and its variants by nsp15. The prominent gel bands indicating specific cleavage are marked by black pentagrams. (**C**) Cleavage of the mL100 and mR100 substrates and their 80-bp variants by nsp15. The approximate sizes of the cleavage products are labeled. (**D**) Identification of the cleavage sites of nsp15 in the Lpca-containing and Rpca-containing short dsRNA substrates labeled with 6-FAM at the 5’ terminus. The three or four strongest cleavage sites in every dsRNA substrate are noted.

After that, we prepared shorter dsRNA substrates to focus on the middle 200-bp region (Figure S2A; Table S1). Here, the corresponding ssRNA for annealing and construction of dsRNA substrates were synthesized utilizing the T7 RNA polymerase S43Y mutant, which was previously found to reduce undesired termination in run-off RNA synthesis and produce RNA with higher terminal homogeneity (Wu et al., 2021). Specific cleavage by nsp15 was observed for the m200 (entire middle 200 bp as described above), mL100 (left half of m200), and mR100 (right half of m200) substrates, but not the m100 (middle 100 bp of m200) substrate (Figure S2B). These results suggested that the preferred cleavage sites for nsp15 are located in both the 386–485-bp (mL100) and 486–585-bp (mR100) regions of the O967 substrate. The cleavage of the mL100 substrate produced two sets of fragments, approximately 60 bp and 40 bp in size. Similarly, cleavage of the mR100 substrate yielded two sets of fragments, approximately 55 bp and 45 bp in size (Figure 2C). The sizes of the cleaved fragments indicated two possible cleavage sites in each substrate. To further locate the cleavage sites for nsp15, we prepared dsRNA substrates that corresponded to 80 bp on the left and 80 bp on the right of the mL100 and mR100 substrates (Figure 2A; Table S1), respectively. After analyzing the sizes of the cleavage products of these substrates (Figure 2C) and comparing them with those from the mL100 and mR100 substrates, we estimated the cleavage sites for nsp15 to be within the 441–450-bp sequence (designated as Lpca, Figure 2A, colored in green) and 526–535-bp sequence (designated as Rpca, Figure 2A, colored in green) of the O967 substrate. To confirm that the originally observed specific cleavage by nsp15 on the O967 substrate occurred in the Lpca and Rpca, we prepared three variants of the O967 substrate with either the Lpca or Rpca or both removed (Figure 2A; Table S1). Deletion of the Lpca or Rpca still maintained the specific cleavage of the O967 substrate by nsp15, while deletion of both abolished the specific cleavage (Figure 2B), confirming that the preferred cleavage sites for nsp15 were indeed located in the Lpca and Rpca.

Interestingly, although the m100 substrate contained both an Lpca and Rpca (Figure S2A), nsp15 did not demonstrate preferential cleavage of this substrate (Figure S2B). Given that both the Lpca and Rpca were close to the end of the m100 substrate, we hypothesized that flanking sequences with adequate length on both sides of the Lpca or Rpca were also required for cleavage by nsp15. We prepared six dsRNA substrates of identical length, all including an Lpca but with various distances from the Lpca to the end of the dsRNA (Figure S2A; Table S1). Results from these substrates showed that as the Lpca approached the end of the dsRNA in either direction, cleavage by nsp15 was weakened (Figure S2C), suggesting that the lengths of flanking sequences on either side of the preferred cleavage sites of nsp15 affected the cleavage efficiency of nsp15.

To locate the exact position of the specific cleavage on the O967 dsRNA substrate by nsp15, we employed short dsRNA substrates labeled with 6-FAM at the 5’ terminus containing the Lpca or Rpca with adequate flanking sequences (Figure 2A). These dsRNA substrates were formed by annealing chemically synthesized RNA oligonucleotides. The cleavage products of these dsRNA substrates by nsp15 were analyzed by denaturing PAGE, and the precise sizes of the cleavage products were determined using the alkaline hydrolysis products of the corresponding ssRNA substrates as ladders. Three to four sites with the highest cleavage efficiencies on every strand of the dsRNA are indicated in Figures 2A and 2D, signifying that nsp15 cleaved at the 3’-side of uridylate in a stretch of consecutive Us. Interestingly, we found that a ssRNA with the same sequence as one of the two strands of the above dsRNA was cleaved by nsp15 in a completely different manner. Nsp15 cleaved almost all U sites and a few C sites in these ssRNA substrates, although the cleavage efficiencies at these sites varied (Figures S2F and S2G). The cleavage at U and C sites in ssRNA by nsp15 was reported previously (Bhardwaj et al., 2006; Frazier et al., 2021; Frazier et al., 2022). However, the cleavage at consecutive Us in dsRNA observed in this study was much more specific and efficient.

In the above assays, the dsRNA substrates were labeled with 6-FAM at the 5’ terminus. To obtain a full observation of the cleavage by nsp15, the dsRNA substrates containing the Lpca as described above were also labeled with 6-FAM at the 3’ terminus (Figure S2D). The preferred cleavage sites for nsp15 deduced on these substrates were consistent with those identified on the 5’-labeled dsRNA substrates. Although the results based on the 5’-labeled dsRNA substrates demonstrated that the most efficiently cleaved site was U24, the results based on the 3’-labeled dsRNA substrates showed that the most efficiently cleaved site was U25 (Figures 2A, 2D, S2D, and S2E), indicating that sequential cleavages might occur in adjacent U sites. Notably, the alkaline hydrolysis products of the ssRNA substrates labeled with 6-FAM at the 3’ terminus did not form standard ladders in the small-sized (1–6 nt) range (Figure S2E). Therefore, the sizes of the cleavage products from these 3’-labeled dsRNA substrates were assigned based on both the alkaline hydrolysis ladder and the cleavage products of the corresponding ssRNA substrates.

### dsRNA cleavage by SARS-CoV-2 nsp15 is sensitive to AU arrangement

Both the Lpca and Rpca were located in areas highly rich in AU content (indicated by orange dashed boxes in Figure 2A). We refer to such areas exceeding 10 bp in length and containing at most one GC base pair as AU-rich areas. We investigated the effect of the AU-rich areas on nsp15 cleavage by replacing various sequences in the Lpca-containing mL100 substrate with GC-rich sequences (Figure S3A; Table S1). When the sequences out of the AU-rich area were substituted by GC-rich sequences, cleavage by nsp15 was not affected (Figure S3B, top). However, when the AU-rich sequences adjacent to the 10-bp core sequence were replaced with GC-rich sequences, we found that not only was the cleavage efficiency of nsp15 reduced, but also the cleavage sites shifted (Figure S3B, bottom). As judged by the sizes of the cleavage products (Figure S3B, bottom), when the sequences on the left side of the AU-rich area were replaced by GC-rich sequences, the cleavage sites shifted toward the right, and vice versa. When the 10-bp or 20-bp AU-rich sequences containing nsp15 preferred cleavage sites from the mL100 substrate were inserted into two other 100-bp dsRNA substrates, they were cleaved similarly by nsp15 (Figures S3C and S3D; Table S1). Despite that the cleavage sites were all within the 10-bp region (Figure 2A), the 20-bp sequences produced stronger cleavages than the 10-bp sequences (Figure S3D). These results suggested that the AU-rich sequences adjacent to the nsp15 cleavage sites facilitated its cleavage. A more quantitative demonstration of the effect of AU-rich sequences around the cleavage sites was shown by short 6-FAM-labeled dsRNA substrates, the −F-L substrate and its variants (Figure 3A). GC replacement outside of the 20-bp AU-rich area showed no significant effect on cleavage within the 10-bp core sequence. However, GC replacement within the 20-bp AU-rich area and adjacent to the 10-bp core sequence reduced the cleavage of nsp15 by half (Figure 3B).

**Figure 3.**
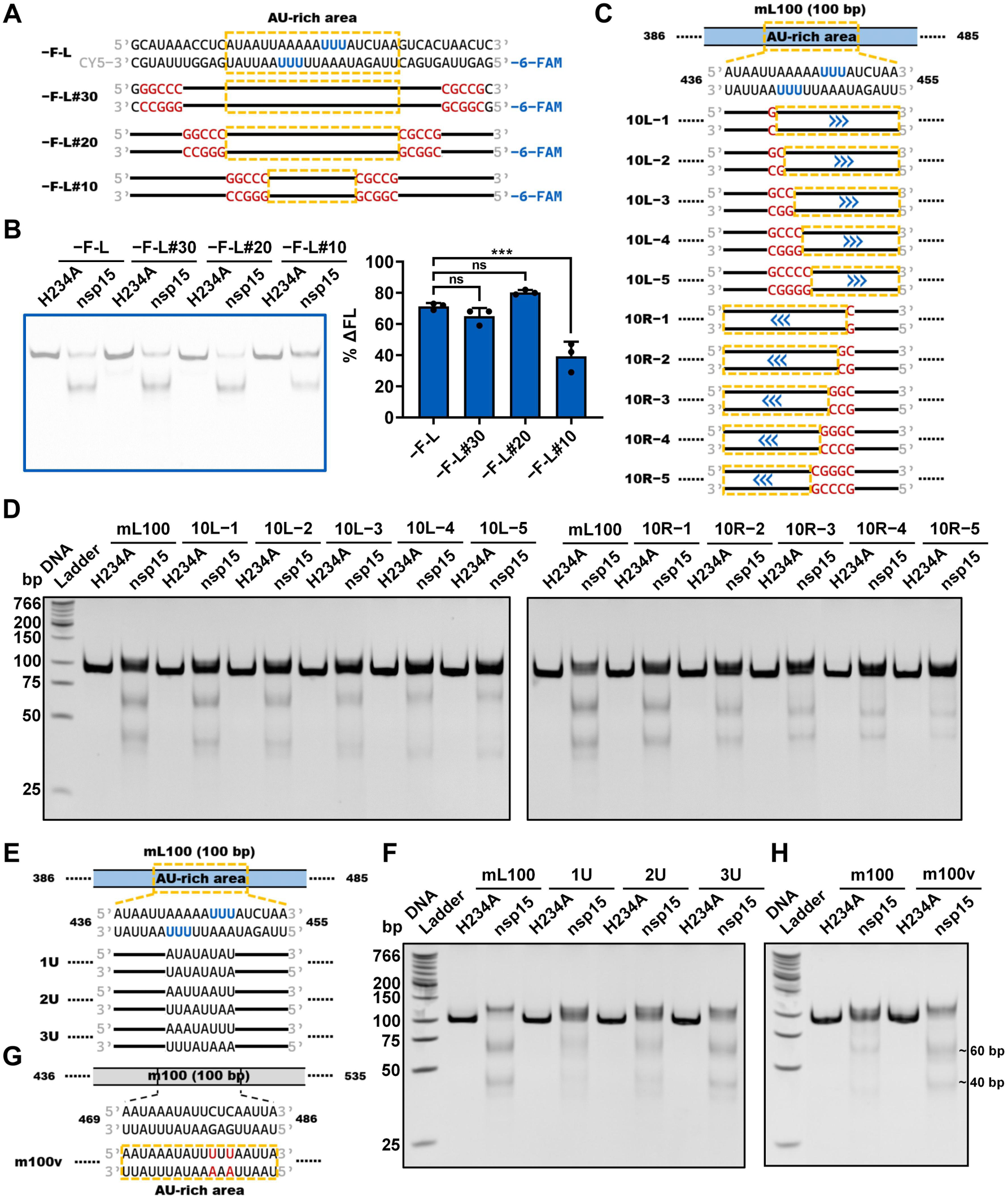
dsRNA cleavage by SARS-CoV-2 nsp15 is sensitive to AU arrangement. (**A**) Schematic representation of the −F-L substrate and its variants used in (**B**). The three U sites with the strongest cleavage in every strand of the −F-L substrate identified previously are shown in blue. The AU-rich areas containing nsp15 preferred cleavage sites are indicated using orange dashed boxes. The GC-rich sequences replacing the original sequences are shown in red. The black lines represent the original sequences. (**B**) Cleavage of the −F-L substrate and its variants as shown in (**A**) by nsp15. Samples were analyzed by native PAGE. The nsp15 H234A mutant was used as a negative control. Reduction of the full-length substrate in every reaction was calculated as % ΔFL. The average and standard deviation of three independent reactions are graphed. Student’s t-test was performed. ns, not significant, p > 0.05; ***p < 0.001. (**C**) Schematic representation of the mL100 substrate and its variants used in (**D**). The blue arrows indicate the shift direction of the dsRNA cleavage sites in the variants as compared with the mL100 substrate. (**D**) Shift of the dsRNA cleavage sites in the substrates shown in (**C**), indicated by the sizes of the cleavage products. An increased distance between the two prominent gel bands indicates a shift to the right, while a decreased distance indicates a shift to the left. (**E**) Schematic representation of the mL100 substrate and its variants used in (**F**). (**F**) Impact of consecutive Us on the cleavage efficiency of nsp15. The concentration of nsp15 or H234A was 10 nM. (**G**) Schematic representation of the m100 substrate and its variant used in (**H**). The two UA base pairs replacing the original CG base pairs are shown in red. The AU-rich area containing consecutive Us ≥3 nt in both strands in the variant is marked with orange dashed boxes. (**H**) Enhancement of nsp15 cleavage by introduction of an AU-rich area containing consecutive Us ≥3 nt in both strands into dsRNA.

We also investigated the effect of GC replacement within the 10-bp core sequence (Figure 3C; Table S1). When the AU sequences on either side were substituted by GC sequences, cleavage by nsp15 decreased, and the cleavage sites were “pushed” toward the other side of the 10-bp core sequence, as judged by the sizes of the cleavage products (Figure 3D). These results suggested that nsp15 preferentially cleaved AU-rich areas in dsRNA.

The observed cleavage sites indicated that nsp15 preferred to cleave consecutive Us rather than individual Us in dsRNA (Figures 2A and 2D). To investigate this property of nsp15, we created the mL100 substrate variants 1U, 2U, and 3U containing no consecutive Us, 2-nt consecutive Us, and 3-nt consecutive Us in the middle of the AU-rich area, respectively (Figure 3E; Table S1). The cleavage of 1U by nsp15 was very weak, while the cleavage efficiency of 3U was significantly higher than that of 2U (Figure 3F). Moreover, stronger cleavage was observed on substrates containing longer consecutive U sequences (Figures S3E and S3F; Table S1). When an AU-rich area containing consecutive Us ≥3 nt in both strands was introduced into another substrate (m100 substrate) by replacing two CG base pairs with UA base pairs (Figure 3G; Table S1), nsp15 cleaved the variant much more efficiently than the original substrate (Figure 3H).

### SARS-CoV-2 nsp15 preferentially cleaves more flexible pyrimidine nucleotides in dsRNA

SARS-CoV-2 nsp15 has been reported to cleave dsRNA via a base-flipping mechanism in two studies (Frazier et al., 2022; Ito et al., 2023). It has also been shown that when ssRNA forms secondary structures, nsp15 prefers to cleave flexible Us located in loops (Bhardwaj et al., 2006; Salukhe et al., 2023). The preference of Us in consecutive AU base pairs in dsRNA by nsp15 may also be due to the flexibility of the Us, as GC base pairs are more thermodynamically stable than AU base pairs. The AU-rich areas in dsRNA may have a more relaxed structure, which may facilitate U-flipping that nsp15 can recognize and cleave. We prepared various 42-bp substrates with different AU distributions or content (Figures 2A and 4A) to examine their thermodynamic stability. The +F-42.1 and −F-42.1 substrates had the same AU content as the +F-L and −F-L substrates, but the AU distribution was more scattered, whereas the +F-42.2 and −F-42.2 substrates had a significantly lower AU content. We analyzed the thermodynamic stability of these substrates using denaturing PAGE (Figure 4B). The results showed that the +F-L and −F-L substrates exhibited the lowest thermodynamic stability and completely denatured in denaturing PAGE. In contrast, the +F-42.2 and −F-42.2 substrates demonstrated the highest thermodynamic stability and resisted the denaturation condition. The +F-42.1 and −F-42.1 substrates exhibited partial denaturation in denaturing PAGE, suggesting intermediate thermodynamic stability. Consistently, nsp15 cleaved the +F-L and −F-L substrates more efficiently than the +F-42.1 and −F-42.1 substrates, and only displayed minimal cleavage of the +F-42.2 and −F-42.2 substrates under identical reaction conditions (Figure 4C), suggesting that low thermodynamic stability of consecutive AU base pairs in dsRNA substrates facilitated cleavage by nsp15.

**Figure 4.**
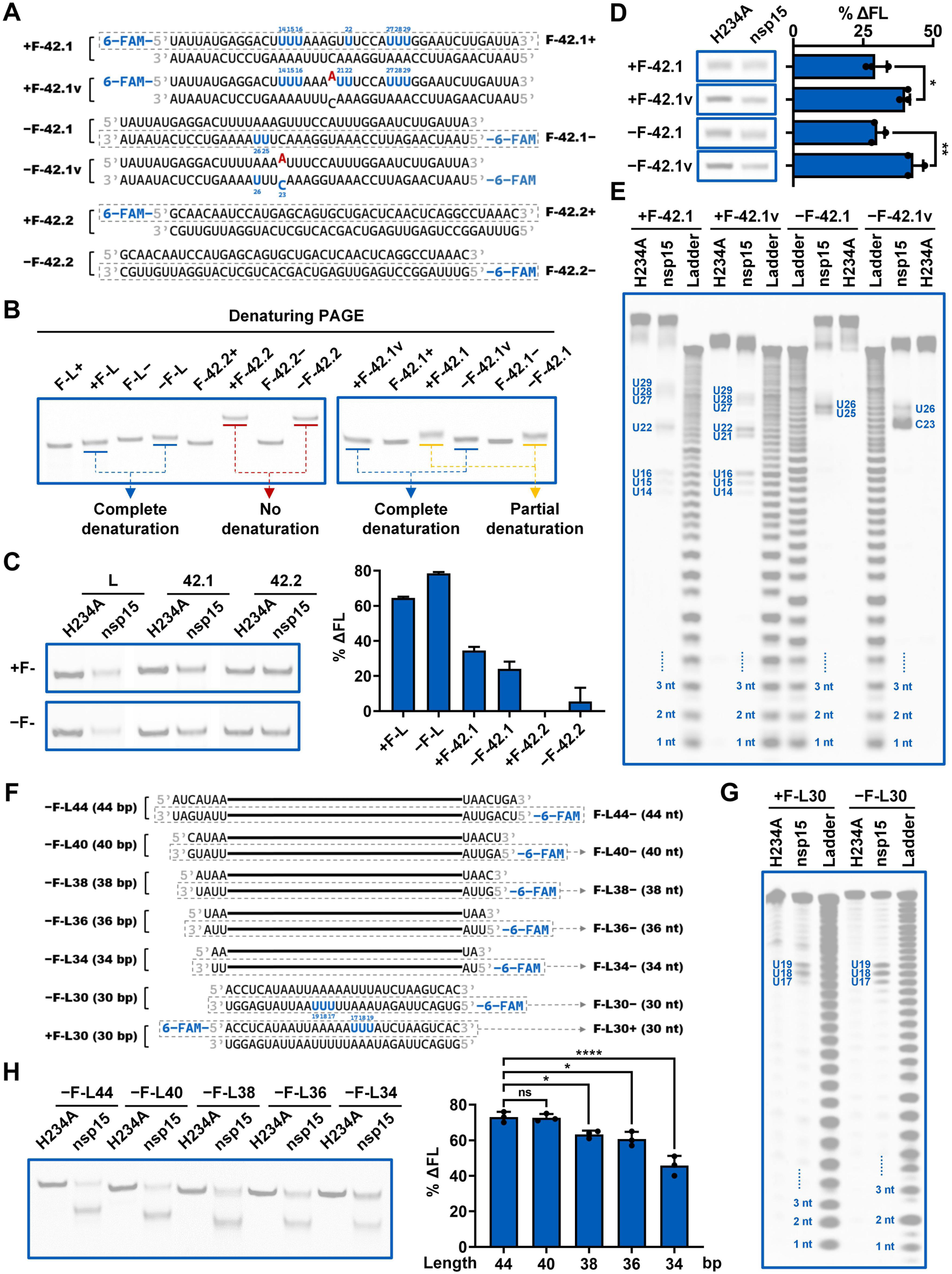
dsRNA thermodynamic stability and length modulate the cleavage of SARS-CoV-2 nsp15. (**A**) Schematic showing the +F-42.1 and −F-42.1 substrates, their variants containing a single base mismatch, and the +F-42.2 and −F-42.2 substrates. The corresponding ssRNA substrates are marked by grey dashed boxes. The preferred cleavage sites of nsp15 on the +F-42.1 and −F-42.1 substrates and their variants are shown in blue. The A base replacing the original G base in the variants of the +F-42.1 and −F-42.1 substrates are shown in red. (**B**) Denaturing PAGE analysis of the indicated RNA substrates. Proximity of the dsRNA gel bands to the corresponding ssRNA gel bands indicates the extent of dsRNA denaturation. (**C**) Cleavage of the indicated RNA substrates by nsp15. Reaction samples were analyzed by native PAGE. (**D**) Cleavage of the +F-42.1, +F-42.1v, −F-42.1, and −F-42.1v substrates showing the impact of a mismatch on nsp15 cleavage. Reaction samples were analyzed by denaturing PAGE. (**E**) Identification of the preferred cleavage sites of nsp15 on the +F-42.1, +F-42.1v, −F-42.1, and −F-42.1v substrates. (**F**) Schematic representation of the variants of the −F-L substrate with various lengths. The three U sites with the strongest cleavage in every strand of the 30-bp variants are shown in blue. The black lines represent the sequence of the −F-L30 substrates. (**G**) Identification of the preferred cleavage sites of nsp15 on the +F-L30 and −F-L30 substrates. The three strongest cleavage sites in every labeled strand are shown. (**H**) Cleavage of the variants of the −F-L substrate with various lengths by nsp15. Reaction samples were analyzed by native PAGE. In (**C, D, and H**), reduction of the full-length dsRNA caused by nsp15 cleavage was quantified as % ΔFL. The nsp15 H234A mutant was used as a negative control. The average and standard deviation for two (**C**) or three (**D and H**) independent reactions are graphed. Student’s t-test was performed in (**D and H**). ns, not significant, p > 0.05; *p < 0.05; **p < 0.01; ****p < 0.0001.

Notably, results obtained on these short RNA substrates revealed that nsp15 cleaved all ssRNA substrates with a low and similar efficiency, unrelated to their AU distribution or content (Figure S4A). In contrast, the cleavage efficiency of dsRNA substrates was highly dependent on their AU distribution and content: the −F-L high AU content dsRNA with more consecutive AU base pairs was cleaved with a considerably higher efficiency than the corresponding ssRNA F-L−; the −F-42.1 high AU content dsRNA with more scattered AU base pairs was cleaved with a similar efficiency compared to the corresponding ssRNA F-42.1−; and the −F-42.2 low AU content dsRNA was barely cleaved compared to the corresponding ssRNA F-42.2− (Figure S4A). At 0.5 mM Mn^2+^ or 5 mM Mn^2+^, the substrate preference of nsp15 on these short RNA substrates was similar (Figure S4A).

To further evaluate the effect of the thermodynamic stability of the dsRNA substrate on its cleavage by nsp15, we created variants of the +F-42.1 and −F-42.1 substrates by replacing a G in the middle with an A to introduce a single-bp AC mismatch (Figure 4A). Compared with the partial denaturing of the original dsRNA, denaturing PAGE analysis of these substrates revealed that the dsRNA substrates harboring the single mismatch were denatured completely, indicating reduction of the thermodynamic stability of dsRNA by the mismatch (Figure 4B). Consistent with our hypothesis, the mismatch enhanced the cleavage efficiency of nsp15 (Figure 4D), and nsp15 showed a strong preference to cleave sites close to the mismatch (Figure 4E). On the +F-42.1 substrate, nsp15 did not cleave the U21 site adjacent to the GC base pair; however, with the introduction of the AC mismatch in the +F-42.1v substrate, nsp15 cleaved the U21 site adjacent to the mismatch position, and cleavage at U22 and U16 sites was also enhanced (Figure 4E). On the −F-42.1 substrate, nsp15 cleaved U26 most efficiently. When the GC base pair was substituted by an AC mismatch in the −F-42.1v substrate, the strongest cleavage switched to C23, right in the mismatch. These results suggested that the mismatch reduced the thermodynamic stability of dsRNA to facilitate base-flipping within the area. It is noteworthy that although A23 was also in the mismatch, no cleavage was observed (Figure 4E), verifying that nsp15 only recognized pyrimidines for cleavage.

### Impact of dsRNA length on cleavage by SARS-CoV-2 nsp15

We previously found that as the Lpca approached the end of the dsRNA, its cleavage by nsp15 diminished (Figure S2C). Additionally, a previous structural study of the nsp15-dsRNA complex reported that dsRNA of approximately 35 bp was sufficient to interact across the nsp15 hexamer (Frazier et al., 2022). Hence, we hypothesized that an adequate length of dsRNA on both sides of the cleavage sites was necessary for binding to nsp15 and for localization of the cleavage sites into the active site of nsp15. We designed variants of the −F-L substrate by lengthening or shortening the flanking sequences on both sides of the cleavage sites (Figure 4F). The substrates +F-L30 and −F-L30 with the 11-bp shortest flanking sequences were still cleaved specifically by nsp15 (Figure 4G). However, the cleavage efficiency increased with an extension of the flanking sequences, and reached a maximum with a 40-bp substrate (Figures 4H and S4B), suggesting that approximately 16-bp flanking sequences at both sides were adequate to provide optimal binding to nsp15. In contrast, the nsp15 cleavage efficiency of the corresponding ssRNA substrates did not vary similarly (Figure S4B), confirming that nsp15 bound to ssRNA or dsRNA using different modes. To examine the disparity in the ability of nsp15 to bind ssRNA and dsRNA, we performed an electrophoretic mobility shift assay (EMSA), using the active-site mutant of nsp15, H234A, which was used in the structural study of the SARS-CoV-2 nsp15 dsRNA complex (Frazier et al., 2022). However, this assay did not reveal significant binding of H234A to either ssRNA or dsRNA (Figure S4C), suggesting that the interactions between nsp15 and both ssRNA and dsRNA were weak or susceptible to interference during electrophoresis.

### SARS-CoV-2 nsp15 is a dsRNA nickase

Although we observed dsRNA breaks resulting from the cleavage by nsp15, the cleavage efficiency for every strand of the dsRNA often varied (Figures 2A, S2D, 5A, and 5H). Moreover, we observed one or two additional bands above the full-length substrate band in the native PAGE analysis of certain cleavage reactions (Figures 5B, S5A, and S5B; Table S1), and these bands were still observed after removal of nsp15 by an RNA purification kit (Figure 5B), indicating that the additional bands did not result from protein binding but instead from dsRNA nicking. We hypothesized that nsp15 may cleave dsRNA in a nicking manner. To test this hypothesis, we created two variants of the 40LR−15 substrate (Figure S3A), in which the consecutive Us were only present on one strand within the AU-rich area (Figure 5C; Table S1). In contrast to the original 40LR−15 substrate, double-strand breaks were not observed for these two variants. Instead, formation of gel bands above the substrate bands were observed (Figure 5D). We also created a variant of the mL100R80 substrate (Figure 2A) containing a pre-existing nick in one strand by annealing three corresponding ssRNA (Figure 5E). We found that this nicked variant mL100R80v was more susceptible to break by nsp15 than the mL100R80 substrate (Figure 5F). Furthermore, the cleavage products of the mL100R80v variant formed a band above the full-length substrate band, indicating that the pre-nicked strand of the variant was further cleaved by nsp15, which was consistent with observations on the cleavage of short dsRNA substrates labeled with 6-FAM (Figures 2A, 2D, S2D, and S2E). All of these results support that nsp15 cleaved dsRNA in a nicking manner. The bands above the full-length substrate band represent the cleavage products in which only one strand is cleaved by nsp15, and the observed dsRNA breaks were due to close nicks in both strands. Despite being an efficient dsRNA nickase, nsp15 was not able to cleave the RNA strand in the RNA-DNA hybrid under identical reaction conditions (Figures 5G and 5H).

**Figure 5.**
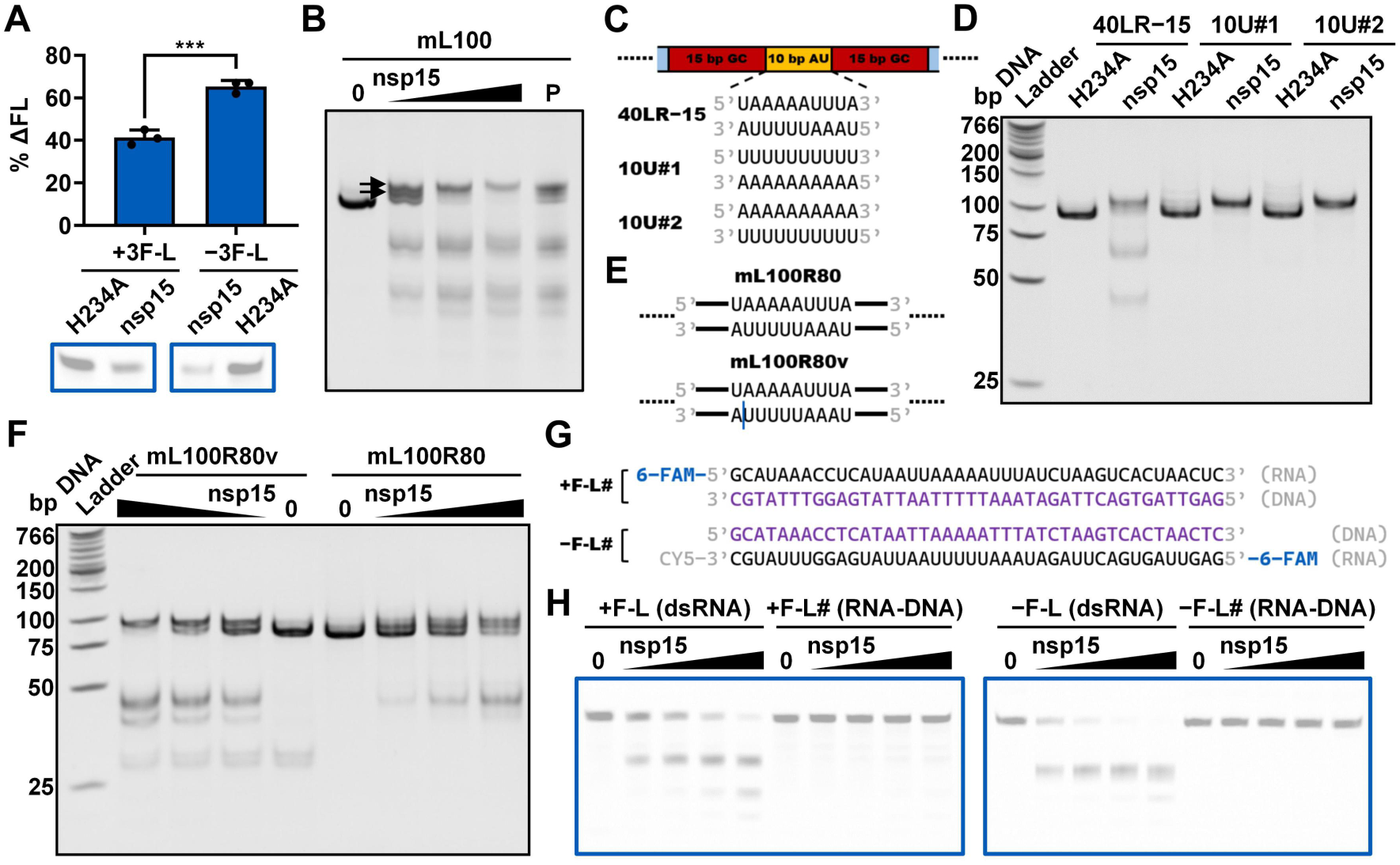
SARS-CoV-2 nsp15 is a dsRNA nickase. (**A**) Cleavage of the 3’-6-FAM-labeled positive-sense or negative-sense strand of the dsRNA substrates shown in **Figure S2D** by nsp15. Reaction samples were analyzed by denaturing PAGE. Reduction of the full-length substrate in every reaction was calculated as % ΔFL. The nsp15 H234A mutant was used as a negative control. The average and standard deviation of three independent reactions are graphed. Student’s t-test was performed. ***p < 0.001. (**B**) Cleavage of the mL100 substrate by nsp15. Reaction samples with varying nsp15 concentrations (0, 5, 10, and 20 nM) were analyzed, along with the RNA purification products (referred to by P) of these samples to exclude protein binding. The gel bands above the full-length substrate gel band are marked by black arrows. (**C**) Schematic representation of the 40LR−15 substrate and its variants in which consecutive Us were only present in one strand within the indicated AU-rich area. (**D**) Cleavage of the 40LR−15 substrate and its variants shown in (**C**) by nsp15. Reaction samples containing the 10-nM H234A mutant or nsp15 were analyzed. (**E**) Schematic representation of the mL100R80 substrate and its nick-containing variant. The nick is marked with a vertical blue line. (**F**) Cleavage of the mL100R80 substrate and its nick-containing variant by nsp15. Reaction samples with varying nsp15 concentrations (0, 2.5, 5, and 10 nM) were analyzed. (**G**) Schematic representation of the RNA-DNA hybrid substrates related to the +F-L and −F-L substrates. The DNA strands are shown in purple. (**H**) Cleavage of the +F-L, +F-L#, −F-L, and −F-L# substrates by nsp15. Reaction samples with varying nsp15 concentrations (0, 5, 10, and 20 nM) were analyzed by denaturing PAGE.

## DISCUSSION

We found that dsRNA length and thermodynamic stability modulated the dsRNA cleavage efficiency of SARS-CoV-2 nsp15 (Figure 4). First, it was important that the dsRNA was of sufficient length to ensure optimal binding to the nsp15 hexamer and to localize the nsp15 preferred cleavage sites to the active site pocket of nsp15. Furthermore, pyrimidines located in relatively thermodynamically unstable areas in dsRNA, such as AU-rich areas and mismatch-containing areas, are more prone to flipping, which allowed for recognition by nsp15 and subsequent cleavage. However, nsp15 accessed most pyrimidines in ssRNA directly without base flipping, which may partially explain the discrepancy between the ssRNA and dsRNA cleavage by nsp15. The thermodynamic stability of ssRNA was mainly determined by its secondary structure, and was not directly correlated with its AU content or AU distribution. Moreover, ssRNA did not exhibit specific binding to the nsp15 hexamer compared to dsRNA. Hence, as the AU content or RNA length decreased, nsp15 exhibited a noticeable decline in cleavage efficiency of dsRNA substrates, but not of the corresponding ssRNA substrates (Figure S4). The inconsistent findings in previous studies on nsp15 substrate preference (Bhardwaj et al., 2004; Frazier et al., 2022; Ivanov et al., 2004) might be due to the different RNA substrates used. We examined the RNA substrates used in these studies and noticed that the only previous study reporting that nsp15 cleaved dsRNA more efficiently (Ivanov et al., 2004) had used a long dsRNA substrate of 1 kb and dsRNA substrate with a single base mismatch. Overall, our study provided sufficient insights into the factors of RNA substrates that determine the substrate preference of coronavirus nsp15.

Coronavirus infection results in the formation of perinuclear double-membrane structures derived from endoplasmic reticulum, including double-membrane vesicles (DMVs), convoluted membranes, and double-membrane spherules, which make up the replication organelles where the viral RTC locates and viral RNA synthesis occurs (V’Kovski et al., 2021). Coronavirus dsRNA, commonly considered as genome replication intermediates, mainly localizes to the DMV interior, a process that is thought to evade detection by cytosolic innate immune sensors (Klein et al., 2020; Knoops et al., 2008; Overby et al., 2010; Wolff et al., 2020). Previous studies reported that coronavirus nsp15 localized with viral RTCs and replicating viral RNA (Athmer et al., 2017; Deng et al., 2017; Heusipp et al., 1997; Shi et al., 1999), suggesting that nsp15 interacted with viral dsRNA intermediates. Furthermore, compared to wild-type virus infection, EndoU-deficient virus infection resulted in increased cytosolic dsRNA (Kindler et al., 2017) or yielded more dsRNA that did not localize with the viral RTC during the early phase (Deng et al., 2017), thus activating host cell dsRNA sensors, such as Mda5, PKR and OAS (Deng et al., 2017; Kindler et al., 2017). This suggests that early during EndoU-deficient virus infection, viral dsRNA intermediates are released to the DMV exterior and detected by cytosolic dsRNA sensors, whereas in wild-type virus infection, nsp15 prevents this from happening using its EndoU activity. Based on our findings, we developed a model showing how nsp15 mediates this process (Figure 6F).

**Figure 6.**
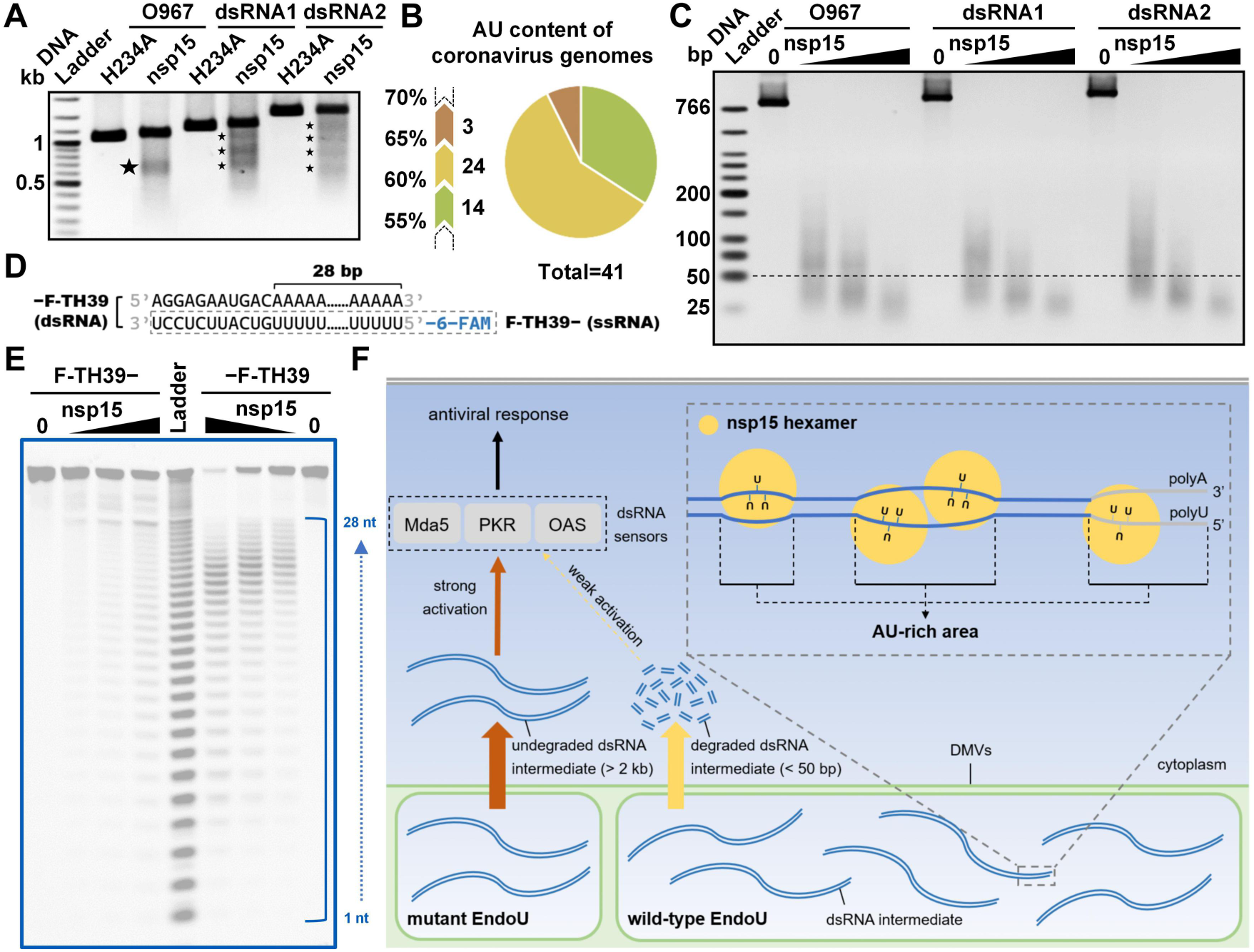
Coronavirus nsp15 degrades viral replication dsRNA intermediates. (**A**) Cleavage of the dsRNA substrates related to the SARS-CoV-2 mini-genome or SARS-CoV-2 S protein gene ORF (dsRNA1 and dsRNA2) by nsp15 with a reaction time of 10 min. The prominent gel bands indicating specific cleavage are marked with black pentagrams. (**B**) Pie diagram grouping the coronavirus genomes based on their AU content. The genomes of 41 known coronavirus species from NCBI were included. Detailed information is listed in Table S4. (**C**) Cleavage of the long (approximately 1 kb) dsRNA substrates mimicking SARS-CoV-2 genome replication intermediates into small fragments by nsp15. Reaction samples with 0.5 mM Mn^2+^ and varying nsp15 concentrations (0, 50, 150, and 450 nM) were analyzed by 2.5% TBE agarose gel electrophoresis. The black dashed line indicates the position of the 50-bp DNA marker. (**D**) Schematic representation of the 39-bp 6-FAM-labeled dsRNA substrate mimicking the dsRNA replication intermediate of the 3’-polyA of the SARS-CoV-2 genome. The ssRNA substrate corresponding to the negative-sense polyU-containing strand is marked by a grey dashed box. (**E**) Cleavage of the polyU-containing ssRNA F-TH39− and dsRNA −F-TH39 substrates by nsp15. Reaction samples with varying nsp15 concentrations (0, 2.5, 5, and 10 nM) were analyzed. The alkaline hydrolysis products of the F-TH39− substrate were used as an RNA size ladder. (**F**) Model depicting how coronavirus nsp15 mediates evasion of host cell dsRNA sensors. Nsp15 preferentially cleaves consecutive Us in AU-rich areas of dsRNA in a nicking manner. The AU-rich areas are widely distributed in coronaviral genome replication dsRNA intermediates. Nsp15 efficiently cleaves dsRNA ≥40 bp, and thus is able to cleave the large dsRNA intermediates into fragments shorter than 50 bp, which may evade cytosolic dsRNA sensors, such as Mda5, PKR, and OAS, even if they are released from the DMV.

Prominent bands were observed in agarose gel electrophoresis analyses of the cleavage products of a high AU content (61%) SARS-CoV-2 mini-genome dsRNA substrate (Figures 1G and S1G-S1K). It was determined that SARS-CoV-2 nsp15 preferentially cleaved consecutive Us in AU-rich areas of dsRNA (Figures 2 and 3) via its dsRNA nickase activity (Figure 5). Upon analyzing the cleavage products of two other dsRNA substrates that were also derived from the genome of SARS-CoV-2 and had a high AU content (64% and 62%), similar prominent bands were observed (Figure 6A) with high cleavage efficiency (Figure 1F). However, when analyzing the cleavage products of a dsRNA substrate derived from the *E. coli* genome and with lower AU content (48%), no prominent bands were observed (Figure S6A), and there was a significantly lower cleavage efficiency (Figure 1F). Interestingly, we found that coronavirus genomes generally have a high AU content, ranging from 55% to 68%, with an average value of 61% (Figure 6B; Table S4). This indicates the potential physiological significance of nsp15 to directly degrade coronaviral dsRNA intermediates. It was reported that OAS 3 selectively binds long dsRNA (>50 bp) (Donovan et al., 2015), PKR prefers to dimerize upon binding to a similar sized dsRNA (>60 bp) (Nallagatla et al., 2011; Robertson and Mathews, 1996), and Mda5 is most efficiently activated by even longer dsRNA (>2 kb) (Kato et al., 2008). Based on the coronavirus genome size of approximately 30,000 nucleotides, unguarded or unprocessed dsRNA intermediates have the ability to strongly activate these dsRNA sensors. Our study showed that nsp15 cleaved dsRNA ≥40 bp with optimal activity (Figure 4H), and we demonstrated that nsp15 was able to cleave long dsRNA with a high AU content into fragments shorter than 50 bp (Figure 6C). Therefore, we hypothesized that after processing by the nsp15 dsRNA nickase, the degradation products of viral dsRNA intermediates are not able to efficiently activate the dsRNA sensors, even if they are released from the DMVs. A previous study reported that the 5’-polyU of the coronavirus negative-sense strand RNA acted as an Mda5-dependent PAMP (Hackbart et al., 2020). Interestingly, in our study, we found that nsp15 did not preferentially target consecutive Us when cleaving ssRNA (Figure S2G). However, we demonstrated that nsp15 cleaved the dsRNA form of a 39-bp 5’-polyU-containing sequence derived from the negative-sense strand of SARS-CoV-2 genome with significantly higher cleavage efficiency compared to cleaving the corresponding ssRNA (Figures 6D and 6E). These results suggested that nsp15 is more likely to remove the 5’-polyU of negative-sense strand RNA by directly cleaving the dsRNA intermediate. Overall, our work supported the mechanism that coronaviruses evade the antiviral response mediated by host cell dsRNA sensors by using nsp15 dsRNA nickase to directly cleave dsRNA intermediates formed during genome replication and transcription (Figure 6F).

## MATERIALS AND METHODS

### Protein expression and purification

DNA fragments encoding wild-type SARS-CoV-2 nsp15 were amplified by PCR from a pET-32a(+) plasmid carrying the nsp15 gene (Ma et al., 2022) and inserted into pET-28a(+) vectors harboring an N-terminal 6× His tag using a ClonExpress II One Step Cloning kit (Vazyme, China) (complete sequence of the nsp15 expression plasmid is listed in Table S3). The constructs were transformed into *E. coli* BL21(DE3)pLysS cells (AngYuBio, China). The cells were cultured in 6 L of LB medium containing 50 µg/mL kanamycin and 34 µg/mL chloramphenicol at 37°C until the OD600 reached approximately 0.8. The flasks containing the culture were then placed at 4°C for 1 h. Then, protein expression was induced by addition of 0.2 mM IPTG, and incubation continued at 16°C for 16 h. The cells were harvested and then stored at −80°C.

Frozen cells were thawed on ice and resuspended in a lysis buffer [50 mM Tris-HCl (pH 8.0 at 25°C), 500 mM NaCl, 5% (v/v) glycerol, 10 mM 2-mercaptoethanol, 10 mM imidazole, and 1 mg/mL lysozyme], then lysed by ultrasonication. The supernatant was collected after centrifugation at 14,000 rpm and 4°C for 1 h and filtered with 0.45-μm filters. The filtered supernatant was loaded onto an Ni-NTA agarose column (Qiagen, Germany) pre-equilibrated with five volumes of the lysis buffer without 2-mercaptoethanol or lysozyme. The column was first washed with 10 volumes of an elution buffer [20 mM Tris-HCl (pH 8.0 at 25°C), 500 mM NaCl, 5% (v/v) glycerol, 10 mM 2-mercaptoethanol, and the indicated concentration of imidazole] containing 40 mM imidazole, and then five volumes of an elution buffer containing 50 mM, 60 mM, and 70 mM imidazole. Most of the His-tagged nsp15 was eluted by an elution buffer containing 200 mM imidazole. The collected eluates were concentrated with a Amicon Ultra-15 30K centrifugal filter (Millipore, USA) and then loaded onto a Superdex 200 column (GE Healthcare, USA) pre-equilibrated with a gel filtration buffer [20 mM Tris-HCl (pH 8.0 at 25°C), 500 mM NaCl, 5% (v/v) glycerol, and 1 mM DTT] for gel filtration chromatography. Fractions containing pure nsp15 were concentrated. Finally, nsp15 was dialyzed against a storage buffer containing 50 mM Tris-HCl (pH 8.0 at 25°C), 100 mM NaCl, 1 mM DTT, 0.1 mM EDTA, 0.1% (v/v) Triton X-100, and 50% (v/v) glycerol, and then stored at −20°C.

A single active site mutation (H234A) was introduced using the Gibson assembly method (primers for cloning are listed in Table S3). The H234A mutant was expressed and purified with the same procedure as detailed above.

The VSW-3 RNA polymerase and the T7 RNA polymerase S43Y mutant used in IVT were expressed and purified as described previously (Wu et al., 2021; Xia et al., 2022).

Protein concentrations were determined with a Bradford Protein Quantitative kit (Bio-Rad, USA), with bovine serum albumin as a standard. Protein purity was analyzed by SDS-PAGE with Coomassie blue staining. Protein size was determined with Precision Plus Protein Standards (Bio-Rad, USA) in SDS-PAGE analysis.

### RNA substrates preparation

#### For long RNA substrates (>500 nt or bp)

The SARS-CoV-2 genome sequence was obtained from the NCBI (NC_045512.2). The DNA templates were synthesized by GenScript or constructed via the Gibson assembly method (primers for cloning are listed in Table S2) and cloned into a pUC19 vector (sequences of plasmids are listed in Table S2). The transcription templates (sequences are listed in Table S1) containing VSW-3 RNA polymerase promoter for positive-sense and negative-sense strand ssRNA were amplified by a three-step PCR using PrimeSTAR Max DNA Polymerase (Takara, Japan) (primers are listed in Table S1) and then purified with an PCR Cleanup kit (Axygen, USA). For the IVT reaction, 35 ng/μL template DNA was incubated with 1.5 U/μL murine RNase inhibitor (New England Biolabs, USA), 0.2 μM inorganic pyrophosphatase, and 0.15 μM VSW-3 RNA polymerase at 25°C for 12 h in an IVT buffer [40 mM Tris-HCl (pH 8.0 at 25°C), 16 mM MgCl_2_, 5 mM DTT, 2 mM spermidine, and 4 mM each of the four NTPs]. Then, 2 U of DNase I-XT (New England Biolabs, USA) was added into 10 μL of reaction mixture and incubation was extended for 30 min at 37°C to remove the DNA templates. The transcripts were purified with a Monarch RNA Cleanup kit (New England Biolabs, USA). The RNA annealing reaction containing 10 mM Tris-HCl (pH 7.4 at 25°C), 50 mM KCl, 200 ng/μL positive-sense strand ssRNA, and 200 ng/μL negative-sense strand ssRNA was carried out on a PCR instrument with a specific annealing program (1. 90°C, 1 min; 2. 90°C, 5 s, −0.1°C per cycle; 3. GOTO step 2, 650×; 4. 4°C, ∞.). The RNA substrates related to the ORFs of the SARS-CoV-2 S protein gene (NC_045512.2), *E. coli ung* or *malE* genes (NC_000913.3), or *Taq* DNA Pol I gene (J04639.1) (sequences of plasmids containing these genes are listed in Table S2) were prepared with the same procedure as detailed above (transcription template sequences and primers are listed in Table S1).

#### For short RNA substrates (≤200 nt or bp)

The transcription templates (sequences are listed in Table S1) containing T7 RNA polymerase promoter for the positive-sense and negative-sense strand ssRNA were amplified by two-step PCR using PrimeSTAR Max DNA Polymerase and the pUC19 plasmid containing the DNA template of the SARS-CoV-2 mini-genome or *E. coli ung* gene with corresponding primers (listed in Table S1), and then purified with a Monarch PCR & DNA Cleanup kit (New England Biolabs, USA). For IVT reactions, 35 ng/μL template DNA was incubated with 1.5 U/μL murine RNase inhibitor, 0.2 μM inorganic pyrophosphatase, and 0.15 μM T7 RNA polymerase S43Y mutant at 37°C for 1 h in an IVT buffer. Then, 2 U of DNase I-XT was added into 10 μL of the reaction mixture and incubation was extended for 30 min at 37°C to remove the template DNA. The transcripts were purified with a Monarch RNA Cleanup kit. The RNA annealing reaction was carried out as described above.

The Gibson assembly method was employed to construct the template-containing plasmids needed for preparing the transcription templates for the positive-sense and negative-sense strand ssRNA of the variants of the mL100, m100, S100, E100, and mLLL substrates (primers for cloning are listed in Table S2). The RNA substrates were prepared with the same procedure as detailed above (transcription template sequences and primers are listed in Table S1). To prepare the nick-containing variant of the mL100R80 substrate, approximately 25 μM positive-sense strand ssRNA synthesized by IVT was annealed with 25 μM each of the two RNA oligonucleotides (synthesized chemically by GenScript) corresponding to the two segments of the negative-sense strand ssRNA.

#### For short 6-FAM-labeled RNA substrates (≤44 nt or bp)

The 6-FAM-labeled and non-labeled RNA oligonucleotides were synthesized chemically by GenScript. The RNA annealing reaction containing 10 mM Tris-HCl (pH 7.4 at 25°C), 50 mM KCl, 25 μM 6-FAM-labeled RNA oligonucleotides, and 25 μM non-labeled RNA oligonucleotides was carried out on a PCR instrument with the specific annealing program detailed above. For preparing the RNA-DNA hybrid substrates, the non-labeled RNA oligonucleotides were replaced with the corresponding DNA oligonucleotides in the annealing reaction.

### RNA cleavage assays

To investigate the substrate preference of nsp15 using the RNA substrates shown in Figure 1, 600 ng of substrate RNA was incubated with the indicated concentration of nsp15 or the H234A mutant to a final volume of 10 μL in an RNA cleavage buffer [50 mM Tris-HCl (pH 7.4 at 25°C), 140 mM KCl, 1 mM DTT, and indicated concentrations of EDTA, MnCl_2_, MgCl_2_, CaCl_2_, ZnCl_2_, and CuCl_2_]. Then, 40 U of murine RNase inhibitor were added in the indicated reactions. Reactions were performed at 37°C for the indicated time (30 min if not otherwise specified) and then terminated by the addition of 10 μL of 2× RNA loading dye (New England Biolabs, USA). Reaction samples were heated at 85°C for 2 min, immediately placed on ice for 2 min, and then analyzed by 1% TAE agarose gel electrophoresis. The gel was stained with ethidium bromide and the RNA was visualized with a UVsolo Touch system (Analytik Jena, Germany). The image was processed, and grey values of the gel bands corresponding to the full-length RNA substrates were quantified with ImageJ software.

To detect the specific cleavage of dsRNA substrates by nsp15, 300 ng of dsRNA substrates were incubated with the indicated concentration of nsp15 or the H234A mutant (5 nM if not otherwise indicated) to a final volume of 10 μL in an RNA cleavage buffer [50 mM Tris-HCl (pH 7.4 at 25°C), 140 mM KCl, 1 mM DTT, and indicated concentrations of EDTA, MnCl_2_, MgCl_2_, and CaCl_2_ (0.5 mM MnCl_2_ if not otherwise indicated)]. Reactions were performed at 37°C for the indicated time (30 min if not otherwise specified) and then inhibited by adding 2 μL of 6x TriTrack DNA loading dye (Thermo Scientific, USA) and placing the reaction samples on ice. Reaction samples were analyzed by 1% TAE agarose gel electrophoresis (for dsRNA substrate length >500 bp) or 12% TBE PAGE (for dsRNA substrate length ≤200 bp). The gel was stained with ethidium bromide and RNA was visualized with a UVsolo Touch system. Image processing was conducted using ImageJ software.

To identify nsp15 cleavage sites using 6-FAM-labeled RNA substrates, 1 μM substrate RNA was incubated with 5 nM nsp15 or H234A mutant to a final volume of 10 μL in an RNA cleavage buffer [50 mM Tris-HCl (pH 7.4 at 25°C), 140 mM KCl, 1 mM DTT, and 0.5 mM MnCl_2_]. Reactions were performed at 37°C for the indicated time (30 min if not otherwise specified) and then terminated by the addition of 10 μL of 2× RNA loading dye. Reaction samples were heated at 85°C for 2 min, immediately placed on ice for 2 min, and then analyzed by 20% TBE-urea PAGE (8 M urea). To generate RNA size ladders, alkaline hydrolysis of the corresponding 6-FAM-labeled ssRNA substrates at a concentration of 6 μM to a final volume of 10 μL in an alkaline hydrolysis buffer [50 mM sodium carbonate (pH 9.4) and 1 mM EDTA] was performed for 15 min at 90°C and quenched with 10 μL of 2x RNA loading dye. 6-FAM-labeled RNA was visualized with a ChemiScope imager (CLiNX, China) using the Cy2 (Ex470BL, Em525/30F) channel. Image processing was performed using ImageJ software.

To determine the nsp15 cleavage efficiency of various 6-FAM-labeled dsRNA substrates by native PAGE analysis, 1 μM substrate RNA was incubated with 20 nM nsp15 or H234A mutant to a final volume of 10 μL in an RNA cleavage buffer [50 mM Tris-HCl (pH 7.4 at 25°C), 140 mM KCl, 1 mM DTT, and 0.5 mM MnCl_2_]. Reactions were performed at 37°C for 30 min and then inhibited by adding 2 μL of 6x TriTrack DNA loading dye and placing the reaction samples on ice. Reaction samples were analyzed by 20% TBE PAGE. 6-FAM-labeled RNA was visualized with a ChemiScope imager using the Cy2 (Ex470BL, Em525/30F) channel. The image was processed and grey values of the gel bands corresponding to the full-length dsRNA substrates were quantified with ImageJ software.

To determine the nsp15 cleavage efficiency of various 6-FAM-labeled RNA substrates by denaturing PAGE analysis, 1 μM RNA substrates were incubated with (unless otherwise stated) 10 nM nsp15 or H234A mutant to a final volume of 10 μL in an RNA cleavage buffer [50 mM Tris-HCl (pH 7.4 at 25°C), 140 mM KCl, 1 mM DTT, and unless otherwise stated, 0.5 mM MnCl_2_]. Reactions were performed at 37°C for the indicated time (30 min if not otherwise specified) and then terminated by the addition of 10 μL of 2× RNA loading dye. Reaction samples were heated at 85°C for 2 min, immediately placed on ice for 2 min, and then analyzed by 20% TBE-urea PAGE (8 M urea). 6-FAM-labeled RNA was visualized with a ChemiScope imager using the Cy2 (Ex470BL, Em525/30F) channel. The image was processed and grey values of the gel bands corresponding to the full-length RNA substrates were quantified with ImageJ software.

### Determination of dsRNA thermodynamic stability

First, 10 μL of 2x RNA loading dye was added to 10 μL of a sample containing 50 mM Tris-HCl (pH 7.4 at 25°C), 140 mM KCl, 1 mM DTT, 0.5 mM MnCl_2_, and 1 μM 6-FAM-labeled substrate RNA. The sample was heated at 85°C for 2 min, immediately placed on ice for 2 min, and then analyzed by 20% TBE-urea PAGE (8 M urea). 6-FAM-labeled RNA was visualized with a ChemiScope imager using the Cy2 (Ex470BL, Em525/30F) channel. Image processing was performed using ImageJ software.

### Electrophoretic mobility shift assay (EMSA)

For the EMSA, 1 μM (for −3F-L and 3F-L− substrates) or 400 ng (for mL100 and mL100+ substrates) of substrate RNA was incubated with the indicated concentration of the H234A mutant to a final volume of 10 μL in an RNA binding buffer [50 mM Tris-HCl (pH 7.4 at 25°C), 140 mM KCl, 1 mM DTT, and 0.5 mM MnCl_2_] for 10 min at 37°C. Then, 2 μL of 6x non-denaturing loading dye [10 mM Tris-HCl (pH 8.0 at 25°C), 0.03% (w/v) bromophenol blue, 0.03% (w/v) xylene cyanol FF, and 60% (v/v) glycerol] was added into a 10-μL reaction sample. Samples were analyzed by 12% TBE PAGE, 6% TBE PAGE, or 3% TBE agarose gel electrophoresis. RNA was visualized as described above. Image processing was performed using ImageJ software.

## SUPPLEMENTAL INFORMATION

Supplemental information can be found online.

## ACKNOWLEDGEMENTS

We thank Prof. Zhi Liu for providing the pET-32a(+) plasmid carrying the nsp15 gene and all lab members for helpful discussion. This project is funded by the National Natural Science Foundation of China (grant 32150009 to B.Z.), The Feng Fundation, and Fund from Science,Technology and Innovation Commission of Shenzhen Municipality (grant JCYJ20210324115811032 to B.Z.). Funding for open access charge: National Natural Science Foundation of China.

## AUTHOR CONTRIBUTIONS

B.Z. conceived the project. X.L.W. and B.Z. designed the experiments. X.L.W. carried out the experiments. X.L.W. and B.Z. analyzed the data. X.L.W. and B.Z. wrote the manuscript.

