## Supplemental information for "SARS-CoV-2 nsp15 preferentially degrades AU-rich dsRNA via its dsRNA nickase activity"

### Figure S1

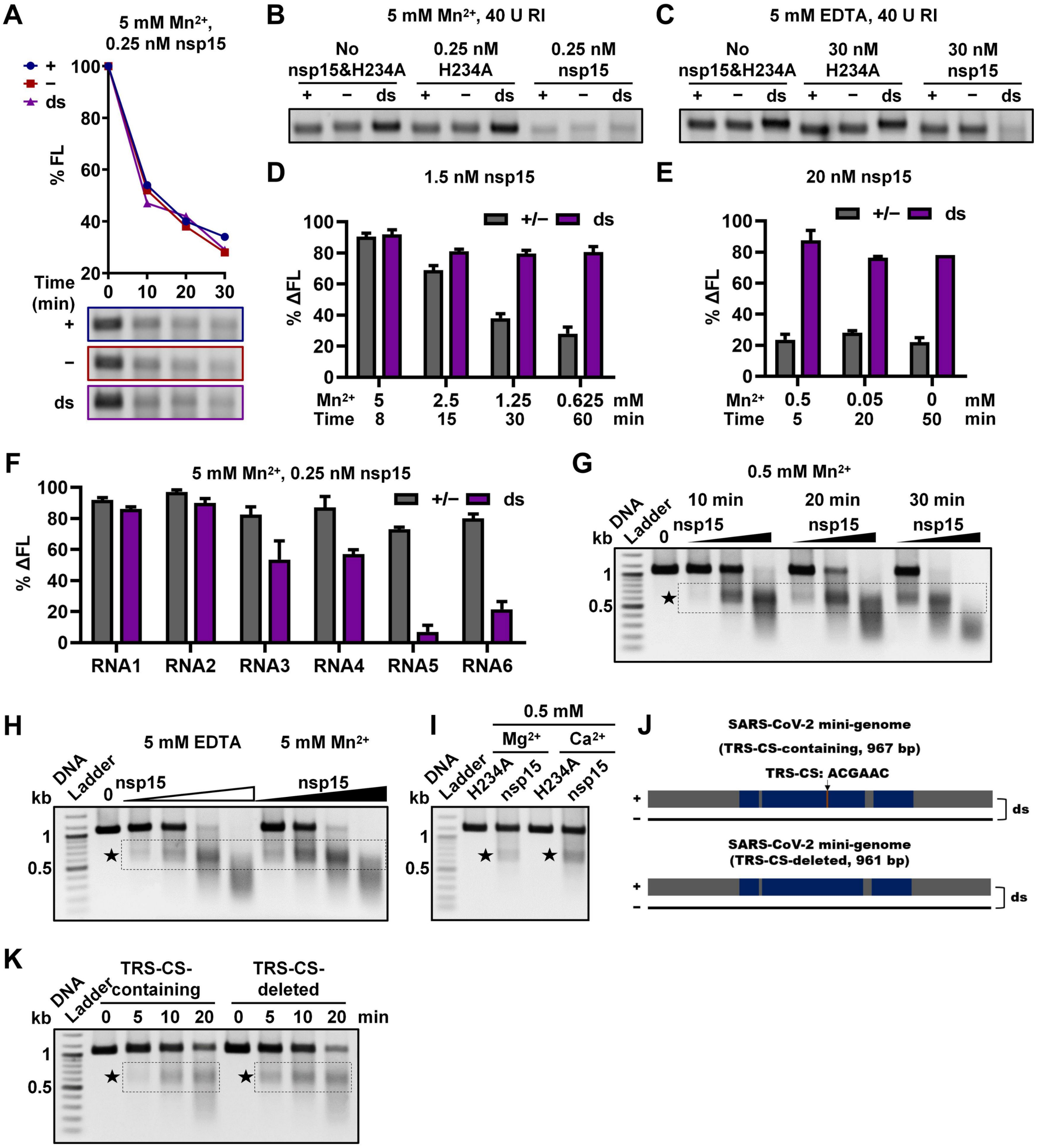

A
5 mM Mn2+, 0.25 nM nsp15
+
−
ds
% FL
Time (min)
0 10 20 30
B
5 mM Mn2+, 40 U RI
No nsp15&H234A
0.25 nM H234A
0.25 nM nsp15
C
5 mM EDTA, 40 U RI
No nsp15&H234A
30 nM H234A
30 nM nsp15
D
1.5 nM nsp15
+/−
ds
% ΔFL
Mn2+ 5 2.5 1.25 0.625 mM
Time 8 15 30 60 min
E
20 nM nsp15
+/−
ds
% ΔFL
Mn2+ 0.5 0.05 0 mM
Time 5 20 50 min
F
5 mM Mn2+, 0.25 nM nsp15
+/−
ds
% ΔFL
RNA1 RNA2 RNA3 RNA4 RNA5 RNA6
G
0.5 mM Mn2+
DNA Ladder
kb
10 min
20 min
30 min
nsp15
H
DNA Ladder
5 mM EDTA
5 mM Mn2+
nsp15
kb
I
0.5 mM
Mg2+
Ca2+
DNA Ladder
H234A
nsp15
kb
J
SARS-CoV-2 mini-genome
(TRS-CS-containing, 967 bp)
TRS-CS: ACGAAC
SARS-CoV-2 mini-genome
(TRS-CS-deleted, 961 bp)
ds
K
DNA Ladder
TRS-CS-containing
TRS-CS-deleted
0 5 10 20 min
kb

**Figure S1. Effect of $Mn^{2+}$ concentration on nsp15 substrate preference. Related to Figure 1.**

(**A**) Cleavage of the ssRNA and dsRNA substrates by nsp15 in the presence of 5 mM $Mn^{2+}$ at various reaction times. (**B**) Cleavage of the ssRNA and dsRNA substrates by nsp15 in the presence of 5 mM $Mn^{2+}$ and 40 U of murine RNase inhibitor (RI). (**C**) Cleavage of the ssRNA and dsRNA substrates by nsp15 in the presence of 5 mM EDTA and 40 U of RI. (**D**) Cleavage of the ssRNA and dsRNA substrates by nsp15 at various $Mn^{2+}$ concentrations (>0.5 mM) and reaction times. (**E**) Cleavage of the ssRNA and dsRNA substrates by nsp15 at various $Mn^{2+}$ concentrations (≤0.5 mM) and reaction times. (**F**) Cleavage of RNA substrates 1–6 by nsp15 in the presence of 5 mM $Mn^{2+}$. (**G**) Cleavage of the dsRNA substrates by various concentrations of nsp15 (0, 1.5, 5, and 15 nM) in the presence of 0.5 mM $Mn^{2+}$ at various reaction times. (**H**) Cleavage of the dsRNA substrates by various concentrations of nsp15 in the presence of 5 mM EDTA or 5 mM $Mn^{2+}$ (0, 15, 30, 60, and 120 nM nsp15 for 5 mM EDTA; 0.125, 0.25, 0.5, and 1 nM nsp15 for 5 mM $Mn^{2+}$). (**I**) Cleavage of the dsRNA substrates by nsp15 in the presence of $Mg^{2+}$ or $Ca^{2+}$. (**J**) Schematic representation of the RNA substrates related to the SARS-CoV-2 mini-genome with TRS-CS deleted. (**K**) Cleavage of the dsRNA substrates with or without TRS-CS by nsp15. All RNA substrates were derived from the SARS-CoV-2 mini-genome unless otherwise indicated. In (**A**), the remaining full-length RNA substrates after nsp15 cleavage were quantified as % FL. In (**D–F**), reduction of the full-length RNA substrates by nsp15 cleavage was quantified as % ΔFL. Reactions containing the same number of nsp15 H234A mutants were used as negative controls. The average and standard deviation for at least two independent reactions are graphed. In (**G–I and K**), the prominent gel bands indicating specific cleavage are marked by a black pentagram and a black dashed box.

### Figure S2

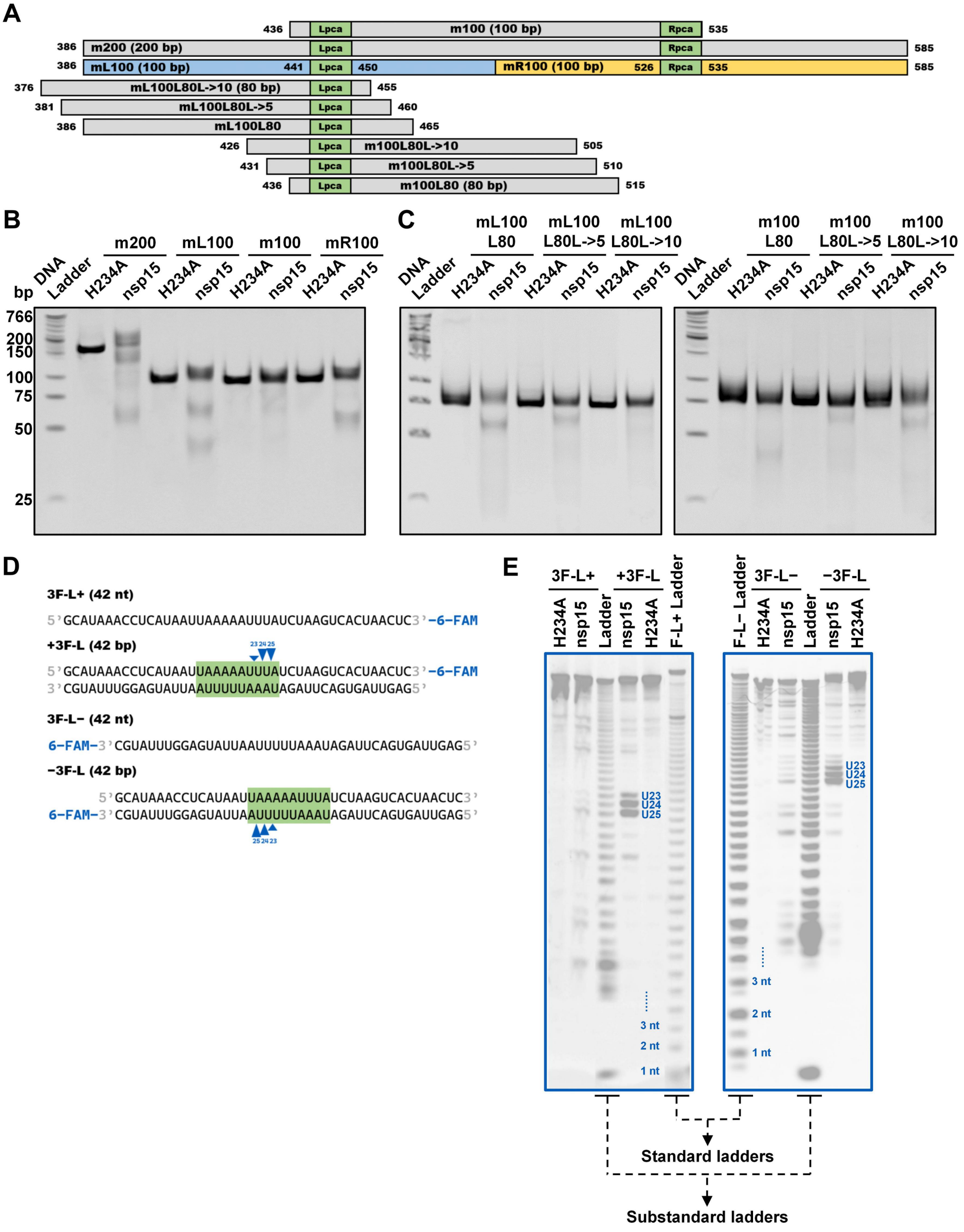
A
436 Lpca m100 (100 bp) Rpca 535
386 m200 (200 bp) Lpca Rpca 585
386 mL100 (100 bp) 441 Lpca 450 mR100 (100 bp) 526 Rpca 535 585
376 mL100L80L->10 (80 bp) Lpca 455
381 mL100L80L->5 Lpca 460
386 mL100L80 Lpca 465
426 Lpca m100L80L->10 505
431 Lpca m100L80L->5 510
436 Lpca m100L80 (80 bp) 515
B
m200 mL100 m100 mR100
DNA Ladder H234A nsp15 H234A nsp15 H234A nsp15 H234A nsp15
bp
766
200
150
100
75
50
25
C
mL100 L80 mL100 L80L->5 mL100 L80L->10 m100 L80 m100 L80L->5 m100 L80L->10
DNA Ladder H234A nsp15 H234A nsp15 H234A nsp15 DNA Ladder H234A nsp15 H234A nsp15 H234A nsp15
D
3F-L+ (42 nt)
5'GCAUAAACCUCAUAAUUAAAAAUUUAUCUAAGUCACUAACUC3'-6-FAM
+3F-L (42 bp)
23 24 25
5'GCAUAAACCUCAUAAUUAAAAAUUUAUCUAAGUCACUAACUC3'-6-FAM
3'CGUAUUUGGAGUAUUAAUUUUUAAAUAGAUUCAGUGAUUGAG5'
3F-L− (42 nt)
6-FAM-3'CGUAUUUGGAGUAUUAAUUUUUAAAUAGAUUCAGUGAUUGAG5'
−3F-L (42 bp)
5'GCAUAAACCUCAUAAUUAAAAAUUUAUCUAAGUCACUAACUC3'
6-FAM-3'CGUAUUUGGAGUAUUAAUUUUUAAAUAGAUUCAGUGAUUGAG5'
25 24 23
E
3F-L+ +3F-L
H234A nsp15 Ladder nsp15 H234A F-L+ Ladder
F-L− Ladder
3F-L− −3F-L
H234A nsp15 Ladder nsp15 H234A
U23
U24
U25
3 nt
2 nt
1 nt
Standard ladders
Substandard ladders

**F**

**F-L+ (42 nt)**

6-FAM-5'GCAUAAACCUCAUAAUUAAAAAUUUAUCUAAGUCACUAACUC3'

**F-L− (42 nt)**

3'CGUAUUUGGAGUAUUAAUUUUUAAAUAGAUUCAGUGAUUGAG5'-6-FAM

**F-R+ (40 nt)**

6-FAM-5'GAUUAAACGAACAUGAAAAUUAUUCUUUUCUUGGCACUGA3'

**F-R− (40 nt)**

3'CUAAUUUGCUUGUACUUUUAAUAAGAAAAGAACCGUGACU5'-6-FAM

**G**

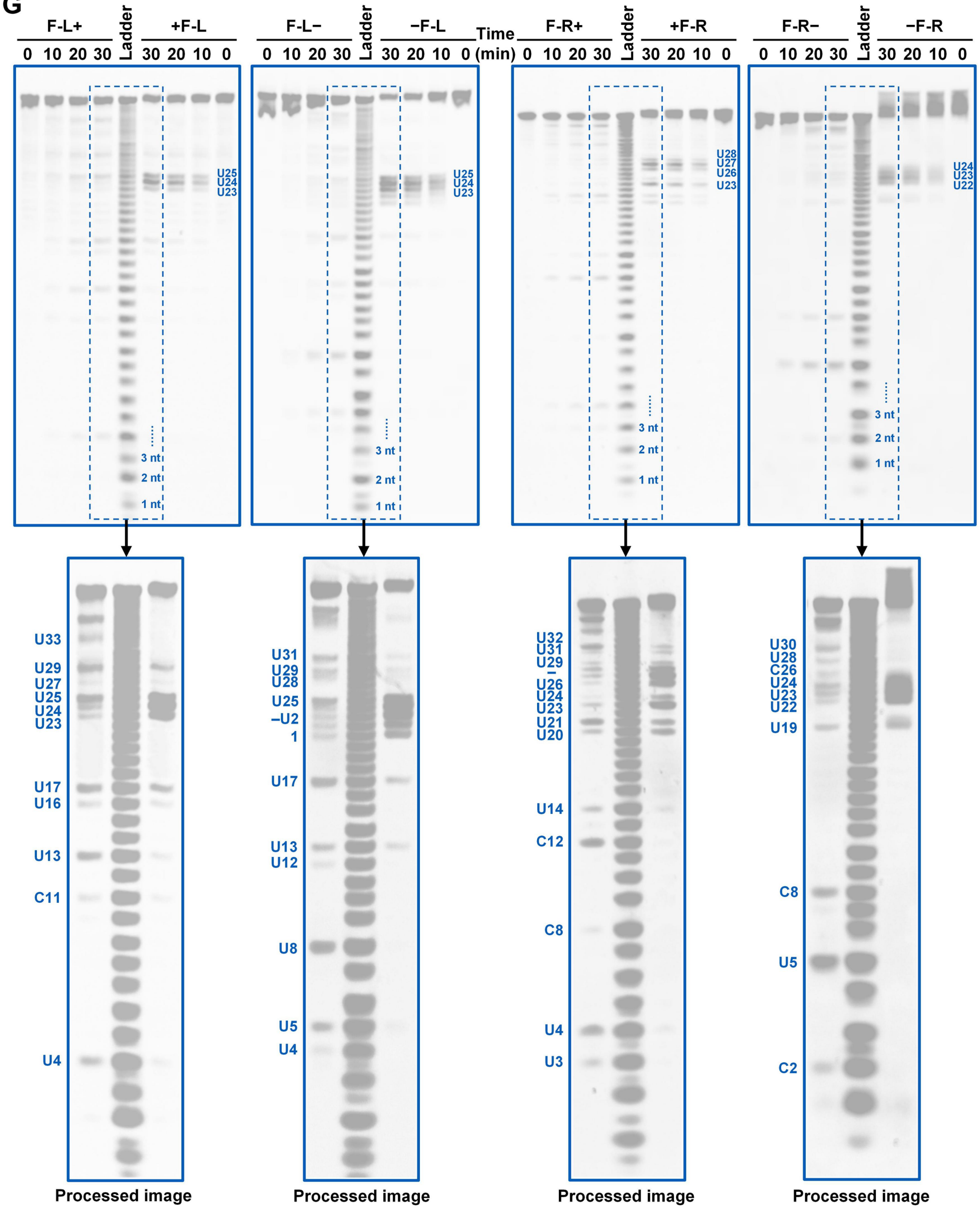

**Figure S2. Cleavage of various dsRNA and ssRNA substrates by nsp15. Related to Figure 2.**

(**A**) Schematic showing the dsRNA substrates used in (**B**) and (**C**). (**B**) Locating the preferred cleavage sites of nsp15 in both 386–485-bp and 486–585-bp regions of the O967 substrate. The cleavage products of the m200, mL100, and mR100 substrates, but not the m100 substrate, formed specific gel bands, indicating that nsp15 preferentially cleaved certain sites in the m200, mL100, and mR100 substrates, but not in the m100 substrate. (**C**) Cleavage of nsp15 on dsRNA containing Lpca at various locations. The cleavage products from the mL100L80, mL100L80L->5, m100L80L->10, and m100L80L->5 substrates but not the mL100L80->10 and m100L80 substrates formed specific gel bands corresponding to cleavages in Lpca, suggesting that nsp15 did not cleave Lpca in the mL100L80->10 and m100L80 substrates. (**D**) Schematic representation of the short Lpca-containing dsRNA substrates labeled with 6-FAM at the 3' terminus and the corresponding 6-FAM-labeled ssRNA substrates. The three sites with the strongest cleavage in every strand of the dsRNA substrates are marked with blue triangles, and the cleavage efficiency is indicated by the height of the triangle. (**E**) Identification of the cleavage sites of nsp15 in the RNA substrates shown in (**D**). The alkaline hydrolysis products of the corresponding 3'-labeled and 5'-labeled ssRNA substrates were used as ladders. The three strongest sites in every dsRNA substrate are noted. (**F**) Schematic representation of the 6-FAM-labeled ssRNA substrates corresponding to the Lpca-containing or Rpca-containing dsRNA substrates labeled with 6-FAM in **Figure 2A**. (**G**) Identification of the cleavage sites of nsp15 in the short Lpca-containing or Rpca-containing RNA substrates shown in (**F**) and **Figure 2A**. The three or four strongest cleavage sites in every dsRNA substrate and most distinguishable cleavage sites in every ssRNA substrate are noted. The brightness and contrast of the dashed box areas were enhanced to identify the ssRNA sites that were weakly cleaved by nsp15.

### Figure S3

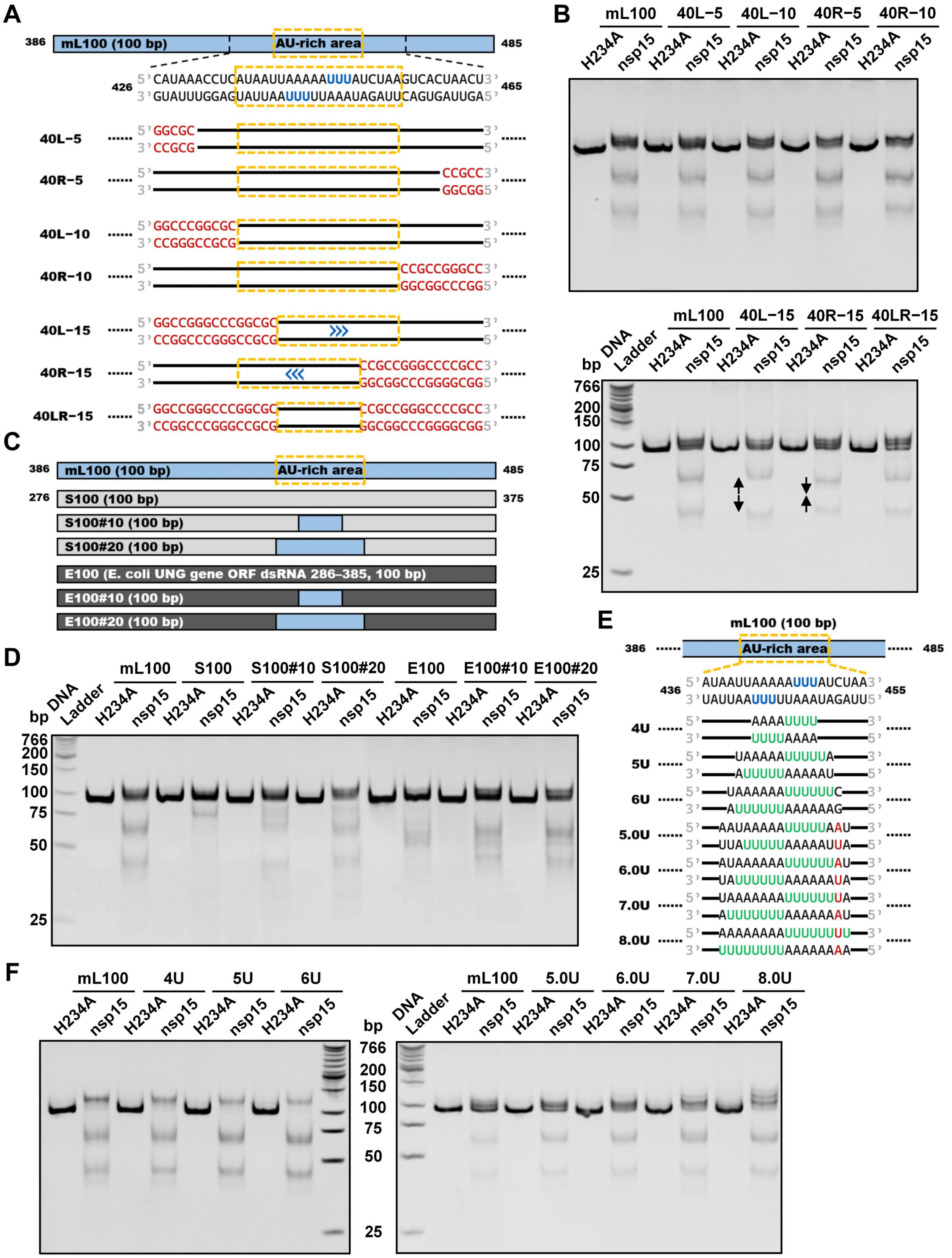
A
mL100 (100 bp)
AU-rich area
386
485
426
465
5' CAUAAACCUCAUAAUUAAAAAUUUAUCUAAGUCACUAACU 3'
3' GUAUUUGGAGUAUUAAUUUUUAAAUAGAUUCAGUGAUUGA 5'
40L-5
5' GGCGC
3' CCGCG
40R-5
CCGCC 3'
GGCGG 5'
40L-10
5' GGCCCGGCGC
3' CCGGGCCGCG
40R-10
CCGCCGGGCC 3'
GGCGGCCCGG 5'
40L-15
5' GGCCGGGCCCGGCGC
3' CCGGCCCGGGCCGCG
40R-15
CCGCCGGGCCCCGCC 3'
GGCGGCCCGGGGCGG 5'
40LR-15
B
mL100
40L-5
40L-10
40R-5
40R-10
H234A
nsp15
40L-15
40R-15
40LR-15
DNA Ladder
bp
766
200
150
100
75
50
25
C
mL100 (100 bp)
S100 (100 bp)
276
375
S100#10 (100 bp)
S100#20 (100 bp)
E100 (E. coli UNG gene ORF dsRNA 286–385, 100 bp)
E100#10 (100 bp)
E100#20 (100 bp)
D
S100
S100#10
S100#20
E100
E100#10
E100#20
E
436
455
5' AUAAUUAAAAAUUUAUCUAA 3'
3' UAUUAAUUUUUAAAUAGAUU 5'
4U
AAAAUUUU
UUUUAAAA
5U
UAAAAAUUUUUA
AUUUUUAAAAAU
6U
UAAAAAAUUUUUUC
AUUUUUUAAAAAAG
5.0U
AAUAAAAAUUUUUAAU
UUAUUUUUAAAAAUUA
6.0U
AUAAAAAAUUUUUUAU
UAUUUUUUAAAAAAUA
7.0U
UAAAAAAAUUUUUUUA
AUUUUUUUAAAAAAAU
8.0U
AAAAAAAAUUUUUUUU
UUUUUUUUAAAAAAAA
F
4U
5U
6U
5.0U
6.0U
7.0U
8.0U

**Figure S3. dsRNA cleavage by SARS-CoV-2 nsp15 is sensitive to AU arrangement. Related to Figure 3.**

(**A**) Schematic representation of the mL100 substrate and its variants used in (**B**). The three U sites with the strongest cleavage on every strand of the mL100 substrate identified previously are shown in blue. The AU-rich areas containing nsp15 preferred cleavage sites are marked with orange dashed boxes. The GC-rich sequences replacing the original sequences are shown in red. The black lines represent the original sequences. The blue arrows indicate the shift direction of the dsRNA cleavage sites in the variants as compared with the mL100 substrate. (**B**) AU-rich sequences flanking the cleavage sites facilitated cleavage by nsp15, while GC substitutions in the flanking sequences resulted in restricted cleavage nearby and a shift of the cleavage sites, as indicated by the size variations between the cleavage products from the mL100 substrate and its variants shown in (**A**). (**C**) Schematic representation of the S100 and E100 substrates and their variants used in (**D**). The sequence of the S100 substrate matched the 276–375-nucleotide sequence of the O967 substrate. The sequence of the E100 substrate matched the 286–385-nucleotide sequence of the *E. coli ung* gene ORF dsRNA substrate shown in **Figure 1E**. The variants contained 10-bp or 20-bp AU-rich sequences from the mL100 substrate. (**D**) The AU-rich sequences from the mL100 substrate conferred nsp15 cleavage on the S100 and E100 substrates. (**E**) Schematic representation of the mL100 substrate and its variants used in (**F**). Consecutive Us are shown in green. The AU or UA base pairs replacing the original CG base pairs are shown in red. (**F**) Consecutive Us enhance the cleavage by nsp15. The concentrations of nsp15 or H234A used for the left and right panel were 10 nM and 2.5 nM, respectively.

### Figure S4

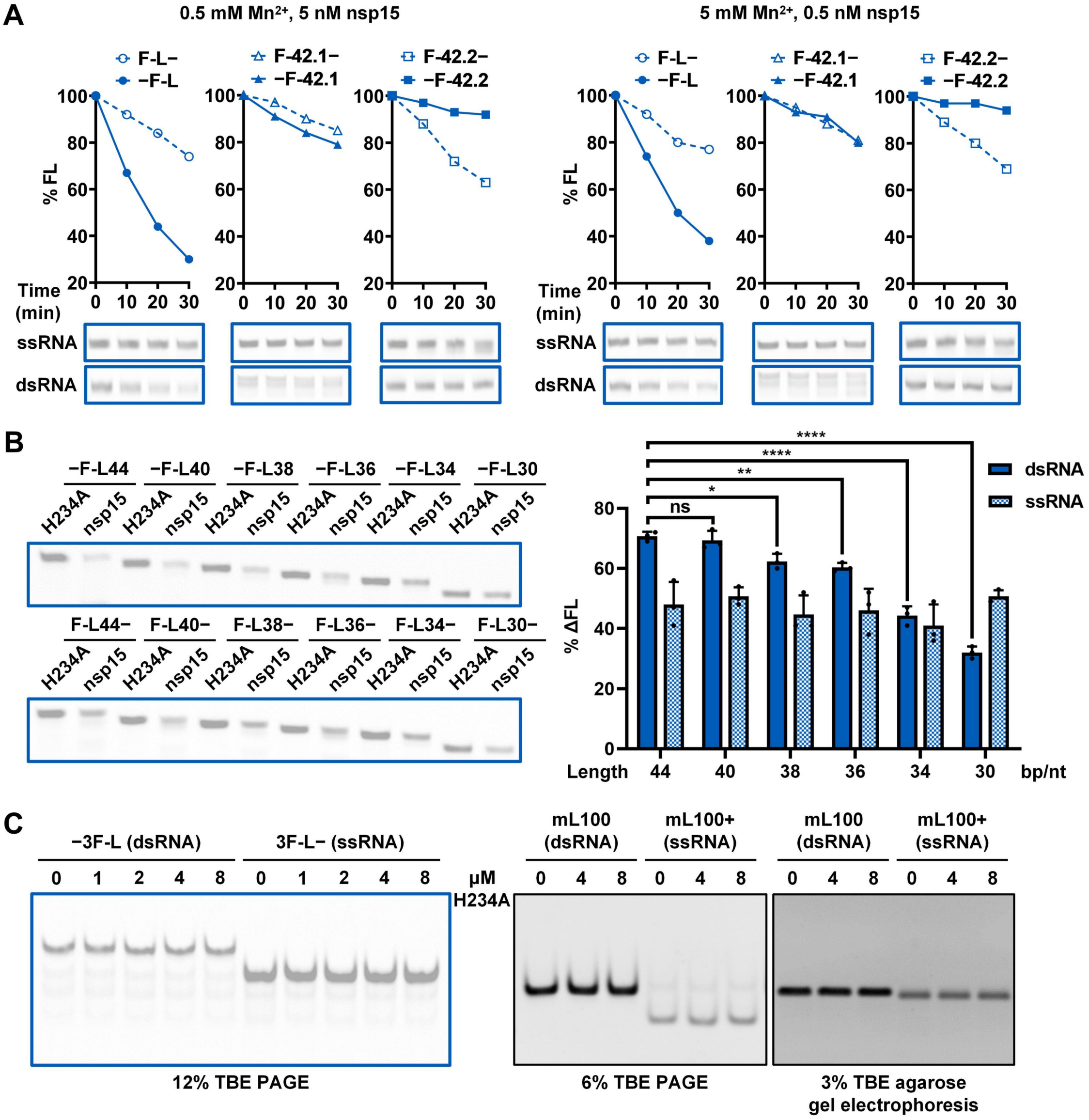
A
0.5 mM Mn2+, 5 nM nsp15
5 mM Mn2+, 0.5 nM nsp15
F-L−
−F-L
F-42.1−
−F-42.1
F-42.2−
−F-42.2
% FL
Time (min)
0 10 20 30
ssRNA
dsRNA
B
−F-L44 −F-L40 −F-L38 −F-L36 −F-L34 −F-L30
F-L44− F-L40− F-L38− F-L36− F-L34− F-L30−
H234A nsp15
dsRNA
ssRNA
% ΔFL
ns
Length 44 40 38 36 34 30 bp/nt
C
−3F-L (dsRNA)
3F-L− (ssRNA)
0 1 2 4 8
μM H234A
mL100 (dsRNA)
mL100+ (ssRNA)
0 4 8
12% TBE PAGE
6% TBE PAGE
3% TBE agarose gel electrophoresis

**Figure S4. Impact of AU content, AU distribution, and RNA length on nsp15 substrate preference. Related to Figure 4.**

(**A**) Cleavage of RNA substrates with various AU content and AU distributions by nsp15 in the presence of 0.5 mM or 5 mM $Mn^{2+}$. Reaction samples were analyzed by denaturing PAGE. Remaining full-length RNA substrates after nsp15 cleavage were quantified as % FL. (**B**) Cleavage of the variants of the −F-L substrate with various lengths and the corresponding ssRNA substrates by nsp15. Reaction samples were analyzed by denaturing PAGE. Reduction of the full-length substrate in every reaction was calculated as % ΔFL. The nsp15 H234A mutant was used as a negative control. The average and standard deviation of three independent reactions are graphed. Student's t-test was performed. ns, not significant, $p > 0.05$; $^{*}p < 0.05$; $^{**}p < 0.01$; $^{****}p < 0.0001$. (**C**) Binding of the nsp15 H234A mutant to dsRNA and ssRNA was detected by electrophoretic mobility shift assay (EMSA). mL100+, the ssRNA substrate corresponding to the positive-sense strand of the mL100 substrate.

#### Figure S5

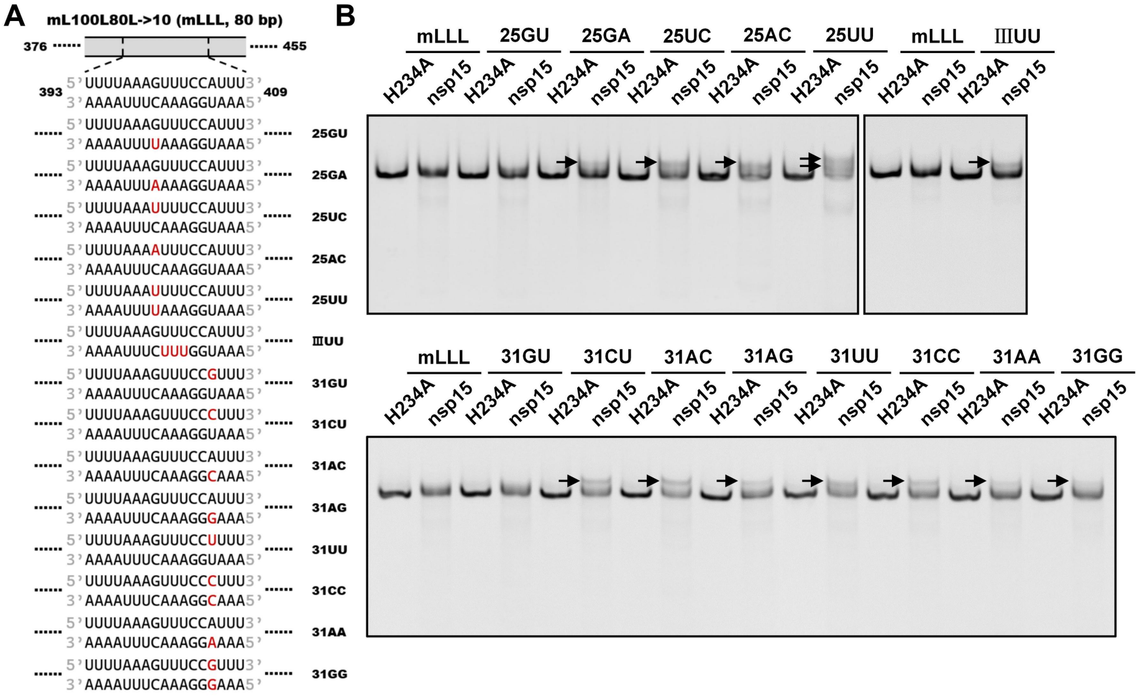


**Figure S5. Cleavage of mismatch-containing dsRNA substrates by nsp15. Related to Figure 5.**

(**A**) Schematic representation of the mL100L80L->10 substrate and its mismatch-containing variants. The bases replacing the original bases in the variants are shown in red. (**B**) Cleavage of the mL100L80L->10 substrate and its mismatch-containing variants shown in (**A**) by nsp15. The gel bands above the full-length substrate gel bands are marked by black arrows.

#### Figure S6

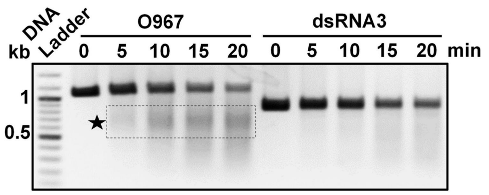


**Figure S6. Cleavage of the dsRNA substrates related to the SARS-CoV-2 mini-genome or *E. coli ung* gene ORF (dsRNA3) by nsp15 at various reaction times. Related to Figure 6.**

The prominent gel bands indicating specific cleavage are indicated by a black pentagram and a black dashed box.
